# Chromosome-level, nanopore-only genome and allele-specific DNA methylation of Pallas’s cat, *Otocolobus manul*

**DOI:** 10.1101/2022.11.30.518596

**Authors:** Nicole Flack, Melissa Drown, Carrie Walls, Jay Pratte, Adam McLain, Christopher Faulk

## Abstract

Pallas’s cat, or the manul cat (*Otocolobus manul*), is a small felid native to the grasslands and steppes of central Asia. Population strongholds in Mongolia and China face growing challenges from climate change, habitat fragmentation, poaching, and other sources. These threats, combined with *O. manul*’s zoo collection popularity and value in evolutionary biology, necessitate improvement of species genomic resources. We used standalone nanopore sequencing to assemble a 2.5 Gb, 61-contig nuclear assembly and 17,097 bp mitogenome for *O. manul*. The primary nuclear assembly had 56x sequencing coverage, a contig N50 of 118 Mb, and a 94.7% BUSCO completeness score for *Carnivora*-specific genes. High genome collinearity within *Felidae* permitted alignment-based scaffolding onto the fishing cat (*Prionailurus viverrinus*) reference genome. Manul contigs spanned all 19 felid chromosomes with an inferred total gap length of less than 400 kilobases. Modified basecalling and variant phasing produced an alternate pseudohaplotype assembly and allele-specific DNA methylation calls; sixty-one differentially methylated regions were identified between haplotypes. Nearest features included classical imprinted genes, non-coding RNAs, and putative novel imprinted loci. The assembled mitogenome successfully resolved existing discordance between *Felinae* nuclear and mtDNA phylogenies. All assembly drafts were generated from 158 Gb of sequence using seven minION flow cells.

## INTRODUCTION

Pallas’s cat (*Otocolobus manul*), or the manul cat, is a small-bodied carnivore native to the montane grassland and shrubland steppe habitats of Central Asia^1^. *O. manul’s* wide geographic range spans varied climatic regions including the Caucuses Mountains, Mongolia, and the Tibetan Plateau, but populations are concentrated at higher altitudes^2,3^. Deemed the “the grumpiest cat in the world” by the BBC documentary series *Frozen Planet II*^4^, manuls are solitary hunters and subsist on a diet of small mammals^5^. Prussian zoologist Peter Simon Pallas assigned the manul cat’s taxonomic name as *Felis manul* in 1776^6^; the genus name *Otocolobus* was proposed in 1842 by Johann Fiedrich von Brandt^7^ and formalized in 1907 based on skull morphology divergence from *Felis*^8^.

Because of its wide home range and low population density, the conservation status of *O. manul* is variable depending on geography. The International Union for Conservation of Nature (IUCN) Red List classifies the species as “Least Concern” as of 2020 after being “Near Threatened” in assessments from 2002 to 2016^1,9^. However, *O. manul* is locally classified as endangered in several countries and either likely or confirmed extinct in some parts of its former range. Populations are also severely geographically fragmented, raising concerns for reduced effective population size and genome heterozygosity^10,11^. The main threats to manul cats are anthropogenic, including poaching, habitat destruction, and the widespread use of rodenticides, which reduce prey populations and result in secondary poisoning of predators^12^. A successful captive breeding program has led to *O. manul* being widespread in zoo collections around the world^13^.

The manul cat is the sole extant species in the monotypic genus *Otocolobus;* phylogenetic analyses surrounding the relationship of this genus to other small-bodied cats are ongoing. A phylogenetic study using select nuclear genes across the felid radiation placed *Otocolobus* as a sister lineage to the genus *Prionailurus*, a taxonomic group that includes the fishing cat (*P. viverrinus*) and leopard cat (*P. bengalensis*) among others^14^. In contrast, a whole-mitogenome phylogeny placed *Otocolobus* closer to *Felis*, which includes the domestic house cat (*F. catus*)^15^. Both studies are concordant with skull morphology data suggesting a Miocene divergence between *Pantheridae* and other cat lineages^16^. Improved genomic resources for the manul cat will increase resolution for phylogenetic analyses within *Felidae*.

High-quality genome assemblies and comparative genomics experiments increasingly include data on the epigenome and repetitive DNA content^17–19^. DNA methylation, the most well-studied epigenetic mark, is critical in embryonic development, genomic imprinting, and X-inactivation with additional influence on gene expression and transposable element (TE) suppression^20–25^. Methyl-cytosines also exhibit a 2-3 fold increase in mutation rate relative to unmethylated cytosines via accelerated deamination^26^, providing a molecular substrate for phenotypic change across evolutionary time. Long-read sequencing permits haplotype phasing based on DNA sequence variants, meaning that DNA methylation can also be segregated by parent of origin^27^. This development is of paramount importance for genome-wide measurement of allele-specific DNA methylation, a signature of imprinted genes. Despite its biological significance, imprinting is chronically understudied due to technical limitations such as bisulfite fragmentation, multi-mapping, and reduced information complexity^28,29^.

Widespread improvements in genome assembly quality reflect reduced sequencing costs, greater local computing power, and improved contiguity via long-read sequencing^30^. Now within reach even for low-resource laboratories, these powerful tools can support conservation efforts for diverse animal species. Here, we provide a high-quality diploid nuclear genome assembly, updated mitogenome, and allele-specific DNA methylation analysis for *O. manul*, all generated exclusively from Oxford Nanopore sequencing reads.

## MATERIALS AND METHODS

### DNA Sample

A five-year-old male manul cat residing at the Utica Zoo (Utica, NY, USA), Tater, was chosen for genome sequencing (Figure 1). Whole blood was collected by jugular venipuncture and shipped frozen, then thawed and combined with 2 volumes of DNA Shield (Zymo Research, Irvine, CA, USA). Genomic DNA was extracted using a Quick DNA Miniprep Plus kit (Zymo Research) yielding 3 μg of DNA from 200 mL of blood. DNA quality was checked using a nano-spectrophotometer (Implen N60, Munich, Germany) and run out on a 1% gel to visualize fragmentation and RNA contamination.

**Figure 1:**
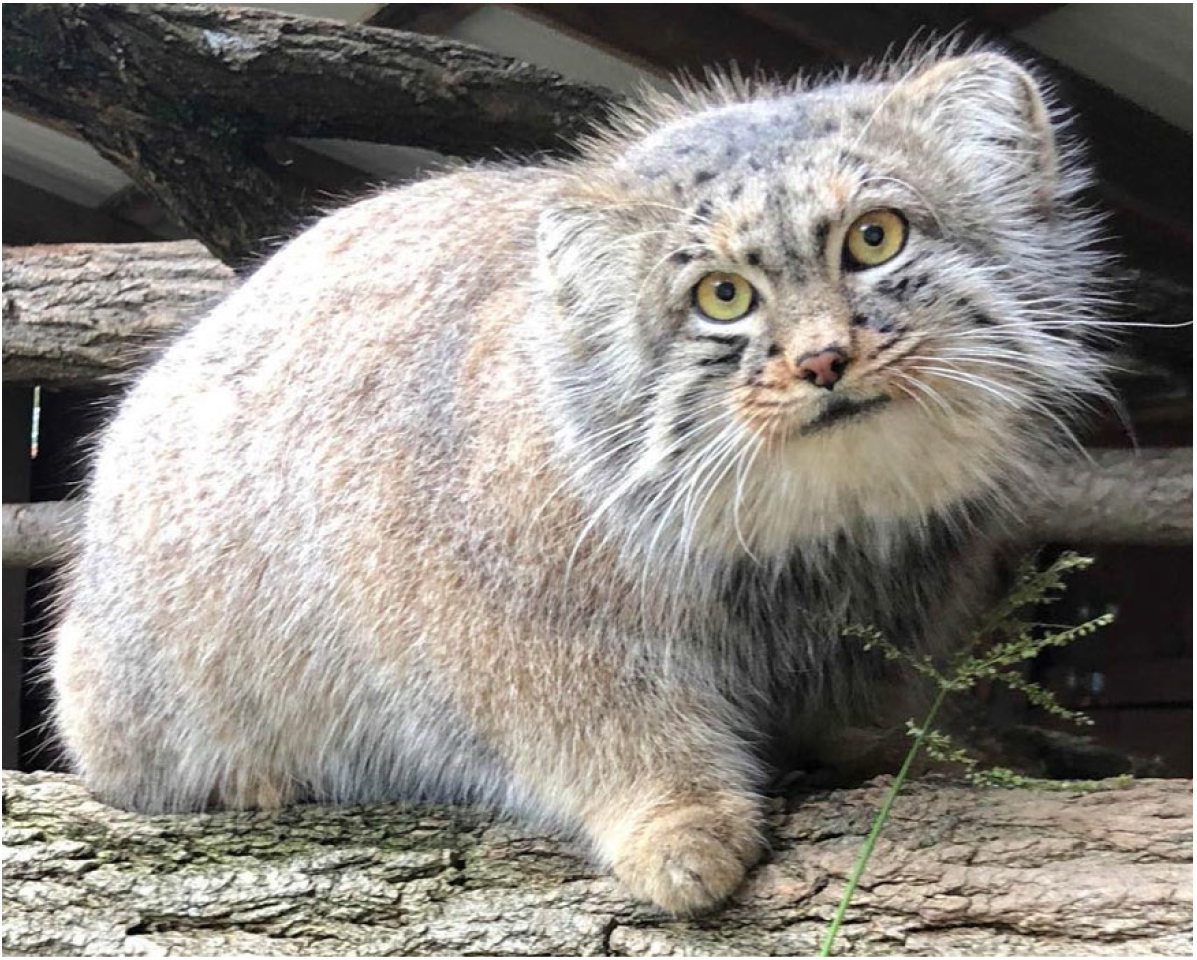
*Otocolobus manul* holotype photo depicting Tater, the five-year-old male manul cat sampled for genome assembly.

### Library Preparation

Seven gDNA library preparations were performed using the SQK-LSK110 kit (Oxford Nanopore Technologies, OX4 4DQ, UK) following a modified version of the manufacturer’s instructions (Nanopore Protocol version GDE_9108_v110_revJ_10Nov2020). In the authors’ modified version, each library prep used 3-5 μg of input DNA and was eluted with 45 μl of elution buffer (EB). Each elution was then split into three 15 μL aliquots to permit one initial flow cell loading step plus two reloads per library prep.

### Sequencing

Sequencing was performed on seven R9.4.1 minION flow cells using Oxford Nanopore’s minKNOW software (v22.05.5) and Guppy basecalling (v6.2.1). A total of three 15 μL DNA aliquots were loaded onto each flow cell with nuclease flushes between reloads per the SQK-LSK110 kit manual’s “Priming and loading” section. Each DNA aliquot was sequenced for 24 hours for a total of 72 hours of sequencing time per flow cell (21 total sequencing days). Guppy was set to fast basecalling during sequencing; post-hoc basecalling was performed using the “super accuracy” model (r941_min_sup_g507). Sequencing summary statistics (Table 1) and a histogram of read length data (Supplemental File 1) are provided.

**Table 1:** *O. manul* nanopore sequencing read summary. Reads were generated from 3 ug of DNA on seven minION flow cells over 21 total sequencing days.

|  |  |
| --- | --- |
| **Number of reads** | 15,349,224 |
| **Number of bases** | 158,248,024,728 |
| **N50 read length (bp)** | 16,601 |
| **Longest read (bp)** | 379,006 |
| **Shortest read (bp)** | 38 |
| **Mean read length (bp)** | 10,309 |
| **Median read length (bp)** | 7,099 |
| **Mean read quality** | 16.26 |
| **Median read quality** | 14.51 |

### Computational Methods

Detailed pipeline information including software versions is provided in Supplemental File 2. The final assembly required the use of cluster computing resources with 2 TB of allocated random access memory (RAM); downstream analyses were performed on two local computers: one running Linux Ubuntu 20.04.1 with 128 GB RAM, an NVIDIA 3090Ti graphics processing unit (GPU), and a 16-core, 32-thread AMD Ryzen 9 5950x central processing unit (CPU), and another running Linux Ubuntu (20.04.1) with 64 GB RAM, two NVIDIA graphics cards (3080Ti and 2080Ti), and a 12-core, 24-thread Ryzen 9 3900x CPU. Abbreviated computational methods are included below.

### Genome Assembly and Polishing

Because there is no single best assembly pipeline for mammalian genomes, we generated draft sequences using a variety of assemblers and polishers. Draft assemblies were generated locally or via institutional cluster computing using Flye^31^, NextDenovo (github.com/Nextomics/NextDenovo), Shasta^32^, and Raven^33^. Polishing was performed with Medaka (github.com/nanoporetech/medaka) for consensus assemblies and Racon^34^ or NextPolish^35^ for non-consensus assemblies. Assembly quality was assessed by searching for *Carnivora*-specific genes with benchmarking universal single-copy orthologs (BUSCO) version 5.3.2^36,37^. The two highest-scoring assemblies from Flye and NextDenovo, respectively, were combined using the Quickmerge metassembler^38^ to increase contiguity while preserving quality. The Purge Haplotigs pipeline was applied for reassignment and removal of allelic contigs^39,40^. Software versions and calls to the assemblers are provided in Supplementary File 2.

### Contamination Detection

Unmapped raw reads were scanned for microbial DNA with Kraken2^41^ and Pavian^42^. We manually inspected the final assembly for spurious contaminant contigs using GC content, unusually high (>1000x) or low (<1x) coverage depth, lack of BUSCO genes, and sequence similarity as metrics. Sequence similarity contamination screening was performed using Megablast versus NCBI’s nucleotide (nt) database^43^. Blobtools2^44^ was used to visualize GC content and read coverage (Supplemental Figure 1). Blobtools also generated the included snail plot. An apparently chimeric 420 kb contig with >1000x coverage and no *Carnivora*-specific BUSCO genes matching *Felidae* by identity, ctg001250, was removed to generate the final primary assembly. Megablast hits to ctg001250 are provided in Supplemental File 3.

### Coverage and Quality Statistics

Assembly coverage was assessed using Mosdepth version 0.3.3^45^. Reference-free, k-mer-based quality statistics were generated with Merqury^46^.

### Repeat Identification

Repetitive DNA masking and classification was performed with RepeatMasker version 4.1.0^47^. Annotation using existing repeat classes, families, and subfamilies was elected based on good representation of *Carnivora* in the Dfam version 3.5 open source repeat library.

### Gene Annotation

GeMoMa v1.8^48,49^ was applied to the primary assembly for homology-based protein prediction with the *Felis catus* gene annotation (Fca126_mat1.0; GCF_018350175.1) used as the reference. Protein models were scored using BUSCO’s protein mode.

### Variant Calling

Variant calling and phasing was performed using the PEPPER-Margin-DeepVariant pipeline^27^. Variant statistics were generated using WhatsHap version 1.4^50^ and VCFtools version 0.1.17^51^. A consensus FASTA for the secondary haplotype was generated by applying all biallelic variants to the final assembly (considered the primary haplotype) with the consensus module of BCFtools version 1.15.1^52^. Runs of homozygosity and genomic demographics were assessed with SMC++^53^.

### DNA Methylation

DNA methylation (5mC) at cytosine guanine dinucleotides (CpGs) was determined by re-basecalling QC-passed FAST5 files with a modified base configuration of GPU-mode Guppy (dna_r9.4.1_450bps_modbases_5mc_cg_sup). The final primary assembly was used as a reference. The resulting modified BAMs (modBAMs) were concatenated together into a single file, then sorted and indexed with Samtools^52^. PEPPER-Margin-DeepVariant was then re-run to generate a haplotagged modBAM; this file and the primary assembly were used as input for ModBAM2BED version 0.6.2 (github.com/epi2me-labs/modbam2bed), which aggregates modified base counts to generate bedMethyl files. These files were then used as input for global and allele-specific methylation analysis.

Allele-specific DNA methylation was analyzed using the R package DSS developer version 2.43.2^54–57^. The package’s two-group statistical comparison module DMLtest was used to identify differentially methylated loci between the two haplotypes. The callDMR module was then used with strict parameters (p.threshold = 0.001, delta = 0.5, minlen = 100, minCG = 15, dis.merge = 1500) to identify multi-CpG differentially methylated regions (DMRs).

To circumvent the relatively low BUSCO quality of our gene annotation, DMR sequences were annotated via lifting of *Felis catus* reference (GCF_018350175.1) gene features onto the *O. manul* assembly using Liftoff^58^. The nearest feature to each DMR was then identified using AGAT (https://github.com/NBISweden/AGAT) and the closest module of BEDTools version 2.30.0^59,60^. DMRs annotated near genes with a “LOC” symbol were additionally annotated with an alias or gene description, when available, via manual look up in NCBI’s Gene database (https://www.ncbi.nlm.nih.gov/gene). Visualizations were generated using MethylArtist version 1.2.3^61^, which required haplotagging the PEPPER output with Longphase^62^, and JBrowse 2^63^ to validate each DMR.

### Mitochondrial Genome

Nanopore reads were aligned to the *Felis catus* mitogenome using Minimap2 (v2.22-r1101)^64,65^. Aligned reads were then downsampled to 10,000 reads using Seqtk version 1.3-r106 (https://github.com/lh3/seqtk) and Flye (v2.9-b1768) in metagenome mode was used for assembly. Mitogenome annotation and manual rearrangement to start the circular mitogenome at COX1 was done as in^66^. A mitogenome phylogeny was built using MUSCLE version 3.8^67^ for alignment and IQ-TREE version 1.6.12^68^ for bootstrapped maximum-likelihood tree inference.

### Scaffolding

The contig-level assembly was scaffolded onto the fishing cat (*P. viverrinus*) reference genome with the scaffold module of RagTag^69^. Alignment of the two genomes was assessed with Dot (github.com/marianattestad/dot), which calls Nucmer^70^. An ideogram of contig positions in the scaffolds was generated with the R pakcage chromoMap (CRAN.R-project.org/package=chromoMap).

## RESULTS

### Sequencing

We isolated DNA for the assembly from the whole blood of Tater, a captive-bred five-year-old male manul cat residing at the Utica Zoo (Utica, NY, USA). A holotype photo is provided (Figure 1). DNA extraction yielded approximately 25 μg of total DNA. DNA was sequenced over 21 days using seven MinION flow cells and base calling was performed using the GPU version of Guppy (v6.2.1). After default quality filtering, sequencing runs generated a total of 15.3 million reads and 158 Gb of sequence with a read N50 of 16.6 kb (Table 1). Of these reads, 66% had a quality score greater than Q20 (i.e., error rate of 1 in 100 base calls) and 20% were greater than Q30 (one error in 1,000 base calls).

### Initial Assembly

We generated multiple *de novo* assemblies and chose the highest-quality result for further analysis based on BUSCO score^37^ (Table 2). Twenty other *Felidae* reference genomes, including three domestic house cat (*Felis catus*) assemblies, provided benchmarking^64,71–81^. Other felid genomes ranged in sequencing coverage from 17x to 159x with an average BUSCO score of 89.56% and a median score of 95.1% (Figure 2); full felid genome assembly statistics used in this comparison are available in Supplemental File 3.

**Figure 2:**
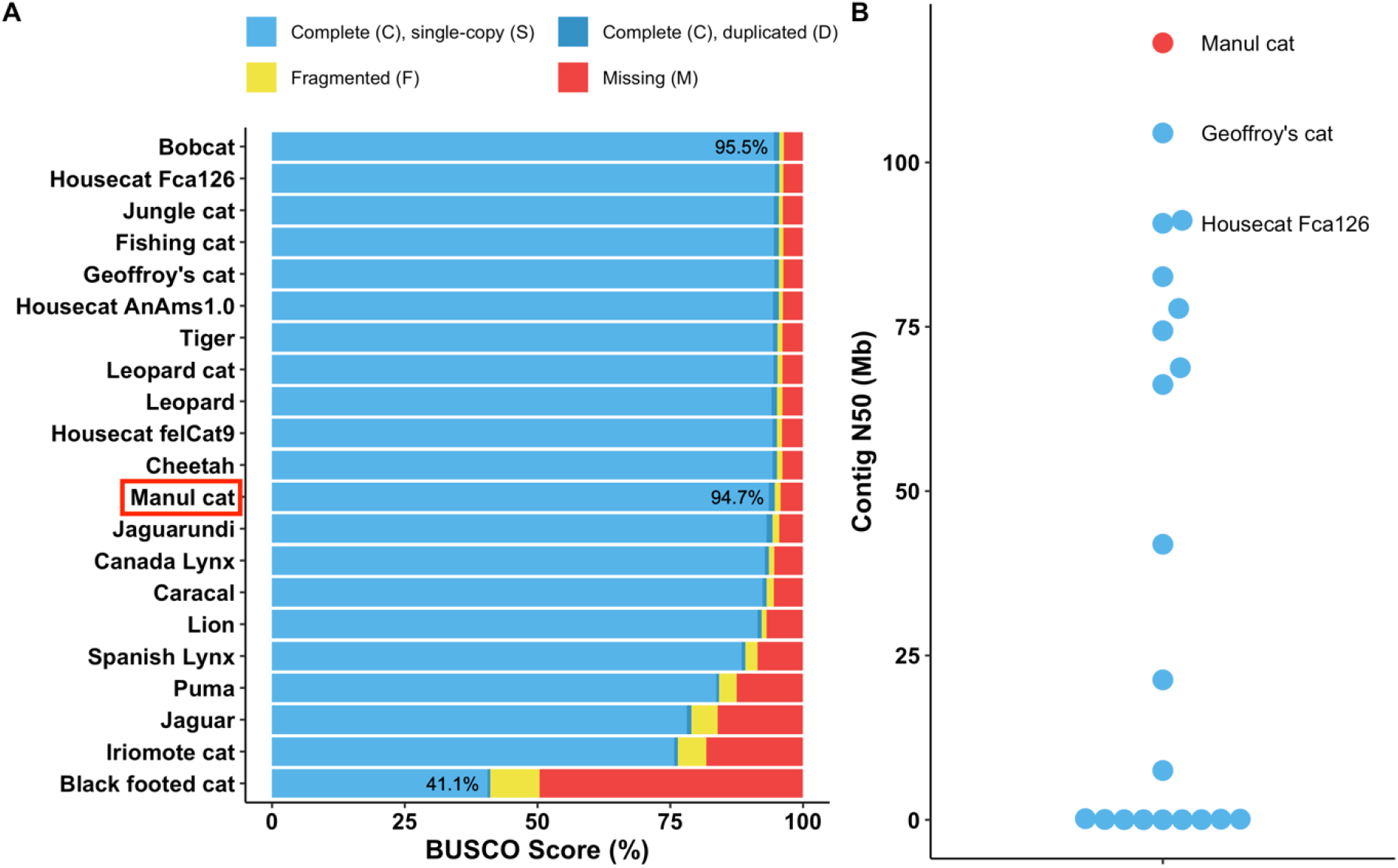
OtoMan1.0 is a near-complete and highly contiguous genome assembly for Pallas’s cat (*Otocolobus manul*). (A) Felid reference assembly completeness was compared to our *O. manul* assembly by locally searching *Carnivora*-specific genes (carnivora_odb10) with BUSCO v5.3.2. OtoMan1.0, a nanopore-only assembly, ranked within 1% of the highest felid assembly BUSCO scores. (B) Contig N50 distribution for *O. manul* and felid reference assemblies. The *O. manul* assembly exhibits high contiguity (N50 = 118.2 Mb) when compared to other *Felidae* genomes, which differ minimally (~ 3 Mb) in size.

**Table 2:** Assembly statistics for selected *O. manul* drafts and reference genomes for fishing cat (*P. viverrinus*) and domestic house cat (*Felis catus*). House cat is the felid species with the most high-quality genome assemblies; fishing cat is more closely related to *O. manul* by nuclear DNA phylogeny^14,15^.

| **Assembly** | **Size (Gb)** | **Contigs/ Scaffolds/ Chromosomes** | **Contig N50 (Mb)** | **BUSCO (% complete)** | **Depth** | **Technique** |
| --- | --- | --- | --- | --- | --- | --- |
| *F. catus* Fca126_mat1.0 (GCF_018350175.1) | 2.42 | 110/71/19 | 90.7 | 95.5 | 76x | PacBio Sequel 2 |
| *P. viverrinus* UM_Priviv_1.0 (GCF_022837055.1) | 2.46 | 255/192/19 | 68.8 | 95.4 | 30x | PacBio Sequel |
| *O. manul* NextDenovo + Medaka | 2.49 | 175/-/- | 36.2 | 94.6 | NA | Nanopore |
| *O. manul* Shasta + Medaka | 2.47 | 714/-/- | 26.3 | 94.1 | NA | Nanopore |
| *O. manul* Flye + Medaka | 2.49 | 1046/-/- | 28.6 | 94.8 | NA | Nanopore |
| *O. manul* Raven + Racon x2 + Medaka | 2.50 | 568/-/- | 9.2 | 94.7 | NA | Nanopore |
| *O. manul* Quickmerge (Flye + NextDenovo) + NextDenovo | 2.49 | 99/-/- | 118.2 | 94.7 | NA | Nanopore |
| *O. manul* Quickmerge + Purge Haplotigs | 2.49 | 62/-/- | 118.2 | 94.7 | NA | Nanopore |
| ***O. manul* Final curated assembly (OtoMan1.0)** | 2.49 | 61/23/19 | 118.2 | 94.7 | 56x | Nanopore |

We ran multiple *de novo* assemblers with relatively low computational memory requirements (i.e., runnable with 64-128 GB local RAM) including Flye, NextDenovo, Shasta, and Raven. Running Flye with the full sequencing read set on a mammal-sized genome required remote high-powered computing resources (≥ 400 GB RAM), so Flye was also run locally with filtered read sets (Supplemental File 3). Filtering reads by length or quality reduced computational resources enough to run assembly software locally but resulted in slightly lower BUSCO scores, even after polishing with the full read set. Therefore, the full QC-passed read set was applied for both drafts used in the final assembly.

The two highest quality draft assemblies were generated by Flye and NextDenovo, both polished once with Medaka (https://github.com/nanoporetech/medaka). Of the two, the Flye sequence achieved the highest BUSCO completeness score for *Carnivora-specific* genes at 94.8%, but was relatively discontiguous with over 1,000 contigs (Table 2). The NextDenovo assembly was more contiguous (175 contigs) but had a lower BUSCO score of 94.6%. Both sequences were appropriate in size for a felid genome at 2.49 Gb. A merged assembly was generated with Quickmerge to preserve the best traits of both drafts; reassignment of allelic contigs was also performed with the Purge Haplotigs pipeline^39,40^, bringing the number of contigs in the best draft assembly down to 62.

### Assembly Curation and Scaffolding

Recent nanopore-only genome assemblies have used anomalous coverage depth and expected GC content as a means of filtering alien contigs^82^. Here, these parameters were assessed using Blobtools2^44^. There was one 420 kb contig with 1,178x average coverage depth in the initial *O. manul* assembly, ctg001250. NCBI megablast matched the sequence to *Felidae*, making a contaminant or endoparasite source unlikely (Supplemental File 3). Examining sequencing read alignment to ctg001250 in JBrowse 2^63^ revealed regions of typical (~60x) coverage interrupted by stretches covered by up to 4,000 reads (Supplemental Figure 2). We concluded that ctg001250 was falsely contiguous based on a low number of reads spanning the high and low coverage areas, perhaps due to misplacement of interspersed repeats. Fifty percent of bases on ctg001250 were masked, and 40% were classified as LINE1 elements by RepeatMasker (Supplemental File 3). BUSCO score was intact following removal of ctg001250 (Table 2).

The final curated assembly, OtoMan1.0, was 2,487,293,883 bp in length with 61 contigs, a contig N50 of 118.2 Mb, and a 94.7% completeness score with BUSCO version 5.3.2 (Figure 2). Total genomic GC content was 41.87%. Comparison of k-mers in both assembly versions (primary and alternate pseudohaplotype) versus the full read set with Merqury^46^ yielded a completeness score of 97.5% and a quality value (QV) score of 45.3. Half of the assembly was covered by 8 contigs (L50) and the largest contig was 218.2 Mb (Supplemental File 3); chromosome A1 in the current reference *F. catus* assembly is 239.4 Mb (RefSeq NC_058368.1). The shortest contig was 133 kb. With local BUSCO v5.3.2 re-scoring of 20 other *Felidae* genomes, a score of 94.7% places our nanopore-only assembly within 1% of the highest quality reference assemblies in this clade, including those generated with short read polishing, chromatin conformation data, and other tools. The assembly’s contig N50 was also the highest among the assessed felid genomes (Figure 2).

OtoMan1.0 was scaffolded onto the high-BUSCO fishing cat (*Prionailurus viverrinus*) reference genome (UM_Priviv_1.0) with the scaffold module of RagTag^69^ (Figure 3). Fishing cat was chosen for alignment-based scaffolding due to (1) a lack of congenial species in *Otocolobus*, (2) the genus sharing its most recent common ancestor with *Prionailurus* rather than *Felis* based on nuclear DNA^14,15^, and (3) the high BUSCO score of the *P. viverrinus* assembly. RagTag joined the 61 *O. manul* contigs into 23 scaffolds with 38 gaps, a total inferred gap length of 371,685 bp, and a chromosome-level N50 of 151.9 Mb (Supplemental File 3). The scaffolds covered all 18 fishing cat autosomes, the X chromosome, and two unplaced BUSCO-containing fishing cat scaffolds (NW_025927612.1 and NW_025927619.1) (Supplemental Figure 3). Two *O. manul* contigs were left unplaced. One of the two unplaced contigs was 593 kb in length and had 98.5% identity to a fragment of the 1.9 Mb *F. catus* Y chromosome (51% query coverage) when assessed with NCBI Megablast^43^; the animal sampled to build UM_Priviv_1.0 was female. The second unplaced *O. manul* contig was 187 kb in length and exhibited high but discontiguous BLAST identity to *F. catus, P. bengalensis*, and other felid sequences.

**Figure 3:**
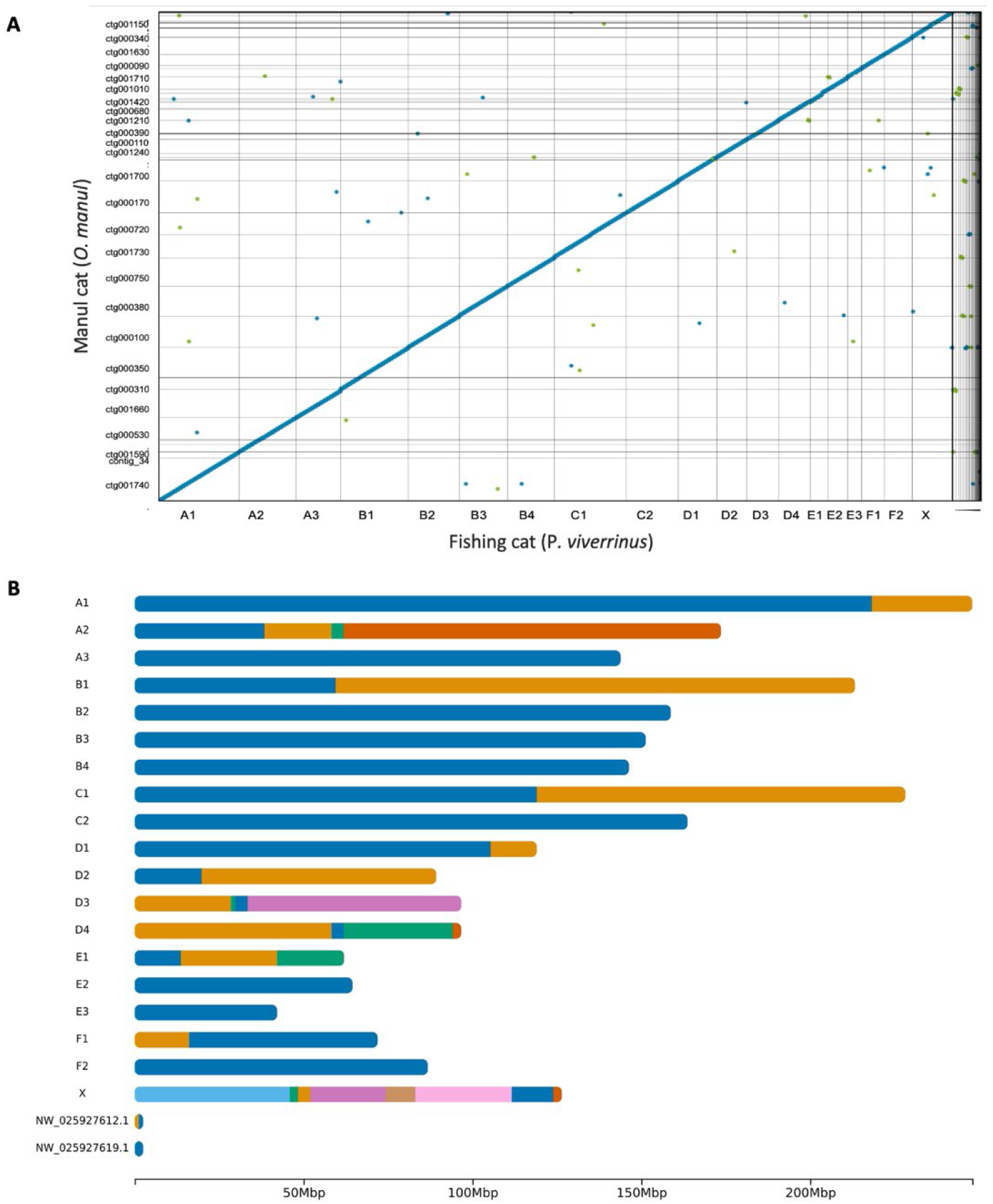
OtoMan1.0 cross-genus alignment and scaffolding. (A) Genome sequence alignment of the fishing cat (*P. viverrinus*) reference genome (UM_Priviv_1.0) and the curated nanopore assembly of *O. manul*, OtoMan1.0, was generated using nucmer^70^ and Dot (https://github.com/marianattestad/dot). The alignment quality reflects gross genome collinearity present across *Felidae*^80,83^. (B) Ideogram of OtoMan1.0 scaffolds generated via alignment to the fishing cat (*P. viverrinus*) reference assembly (UM_Priviv_1.0). Color transitions represent breakpoints between contigs on the scaffold. All 19 felid chromosomes were covered by OtoMan1.0 contigs; contigs were also aligned to two BUSCO-containing fishing cat unplaced scaffolds while two of the 61 *O. manul* contigs were unplaced. Correlations between the fishing cat chromosome lengths and *O. manul* scaffold lengths are provided in Supplemental Figure 3.

### Repetitive DNA

Repetitive DNA, particularly interspersed repeats, are valuable drivers of genetic diversity in vertebrates^84,85^. Since carnivores are well-represented in transposable element (TE) databases, we classified and annotated *O. manul* repetitive DNA with RepeatMasker (Table 3)^47^. Overall, 34.72% of the assembly was classified as repetitive DNA, with a majority of retroelements (26.83%) and fewer DNA elements (2.8%), simple repeats (3.2%), and low-complexity regions (1.4%). These results are consistent with the dominance of retroelements in mammalian genomes^86,87^. Local analysis of other cat assemblies with RepeatMasker indicated that OtoMan1.0’s repeat complement was consistent with other members of *Felidae* (Figure 4). Conservation of global repeat content in this clade was high: the 18 analyzed species varied by 3.67% for total bases masked, 2.72% for LINE content, 0.16% for SINE content, and 0.15% for DNA element content (Supplemental File 3).

**Figure 4:**
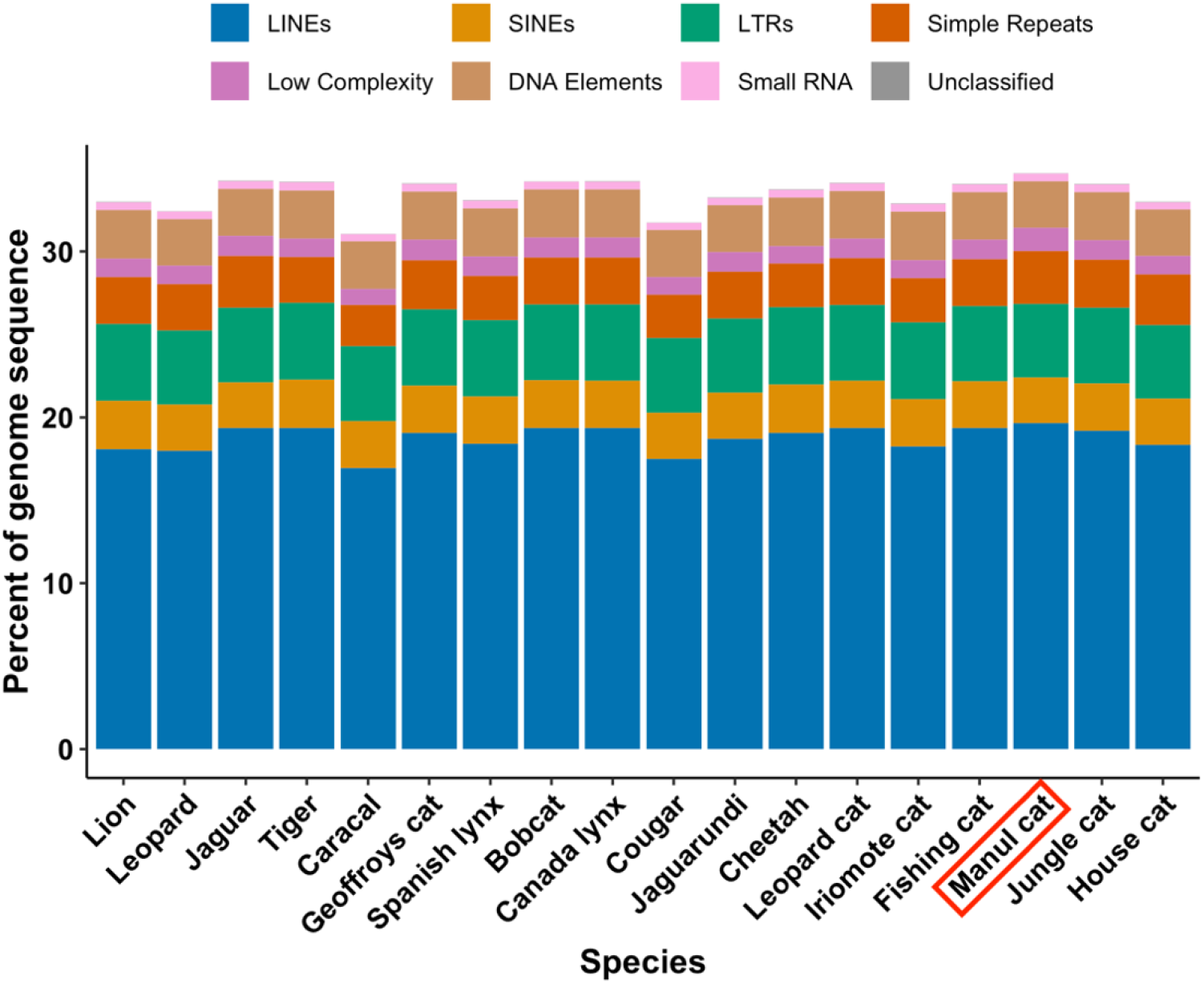
Repetitive element content of the assembled *O. manul* genome and 17 other felid species assessed with RepeatMasker^47^. Manul cat repeat content was highly similar to other members of *Felidae* and dominated by retroelements.

**Table 3:**
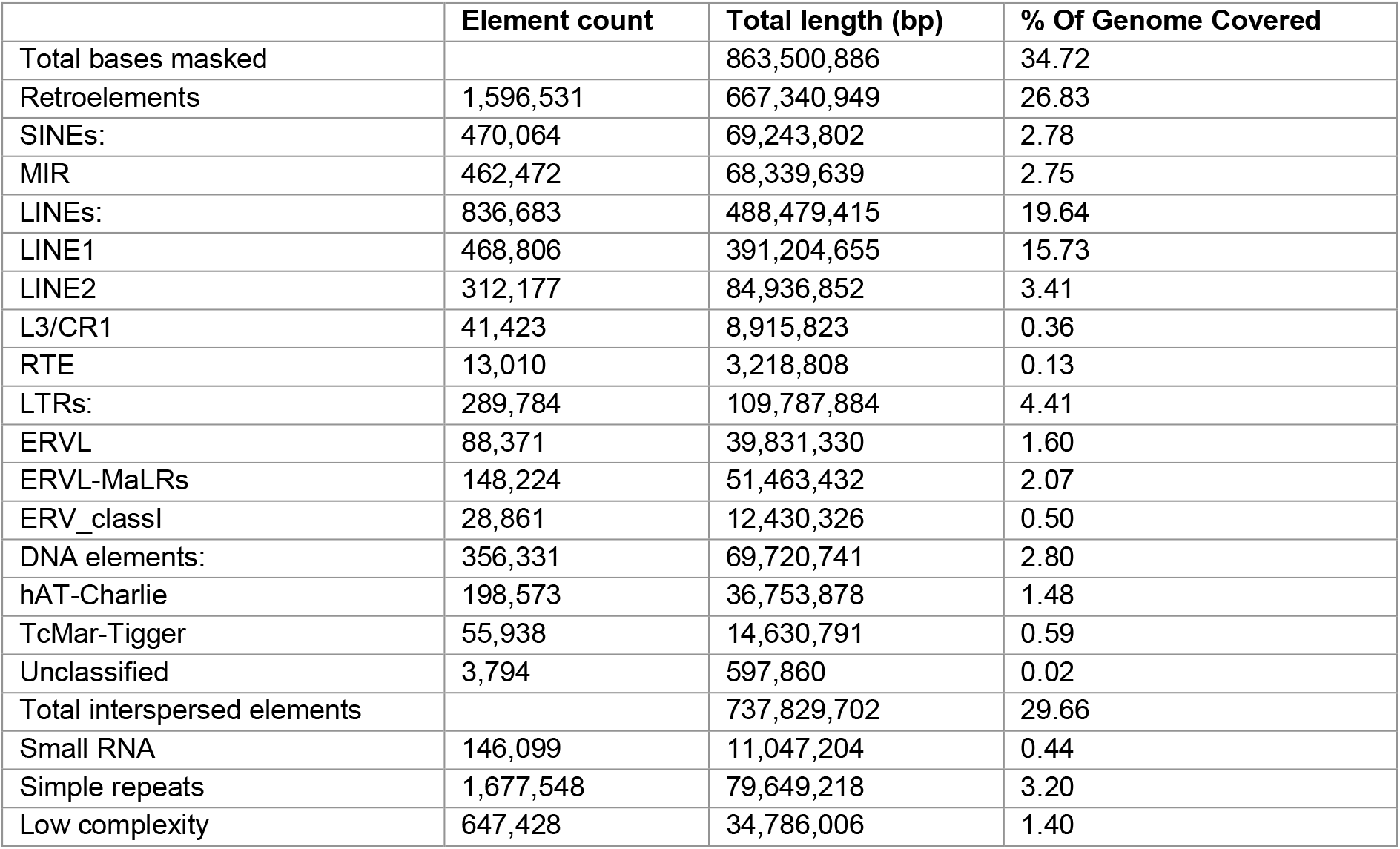
Repetitive DNA classification results for the manul cat (*Otocolobus manul*) genome assembly, OtoMan1.0, generated with RepeatMasker^47^.

### Diploid Genome

After read phasing OtoMan1.0 with the PEPPER-Margin-DeepVariant pipeline^27^, approximately 83% of the assembly was covered by 6,613 phase blocks with a block NG50 of 0.53 Mb (Supplemental File 3). As expected, most variants were single nucleotide variants (SNVs) and small insertions and deletions, with fewer large indels. Of 2,045,611 heterozygous variants detected, 76.3% (1,561,420) were phased and 1,087,364 (53.2%) were phased, biallelic SNVs. The transition-transversion ratio (Ts/Tv) for biallelic SNVs was 2.12, in line with highly methylated mammalian genomes (Supplemental File 4). The software employed for variant calling does not detect inversions.

An alternate pseudohaplotype assembly was generated by switching biallelic variants from the PEPPER variant call file into the primary, contig level assembly using the consensus model of BCFtools^52^. The resulting sequence was 2,484,282,002 bp in length with a contig N50 of 118.0 Mb. The alternate assembly had a 96.8% k-mer completeness score and Q score of 44.8 when assessed with Merqury and contained 94.8% of complete *Carnivora*-specific genes (carnivora_odb10) with BUSCO version 5.3.2. The primary assembly alone was 96.9% complete via Merqury with a Q score of 46.1.

Genome heterozygosity and effective population size for *O. manul* were also assessed using variant calls. Heterozygosity was 0.048% based on the number of called heterozygous SNVs (1,184,174) divided by the total callable bases in the contig-level primary assembly and 0.082% using all 2,045,611 heterozygous variants combined (SNVs and indels). Population size history inferences for *O. manul* based on genomic runs of homozygosity were conducted with SMC++^53^. The results suggested an effective population size (N_e_) near 10,000 with recovery from a bottleneck approximately 3,000 generations ago (Supplemental Figure 4).

### Annotation

Homology-based gene annotation was performed with the Gene Model Mapper (GeMoMa) pipeline version 1.8 using the *F. catus* (GCF_018350175.1) RefSeq annotation and assembly as references. The *F. catus* reference pipeline generated 21,909 amino acid predictions with an 86.2% protein-mode BUSCO score (Supplemental File 3). GeMoMa was also run using the fishing cat (*P. viverrinus*) reference assembly annotation, but BUSCO completeness was slightly lower at 85.3%.

### *Otocolobus manul* genome deposit

Before scaffolding, the complete primary assembly for *O. manul* was 2,487,293,883 bp in length with 61 contigs and a contig N50 of 118,180,270 bp (Figure 5). Mean read coverage depth was 56x. Scaffolding to the fishing cat (*P. viverrinus*) reference genome generated 23 scaffolds covering all 18 felid autosomes and the X chromosome with a scaffold N50 of 151 Mb. The scaffolded primary assembly was deposited as BioProject PRJNA885133 and the contig-level alternate haplotype assembly as BioProject PRJNA889808.

**Figure 5:**
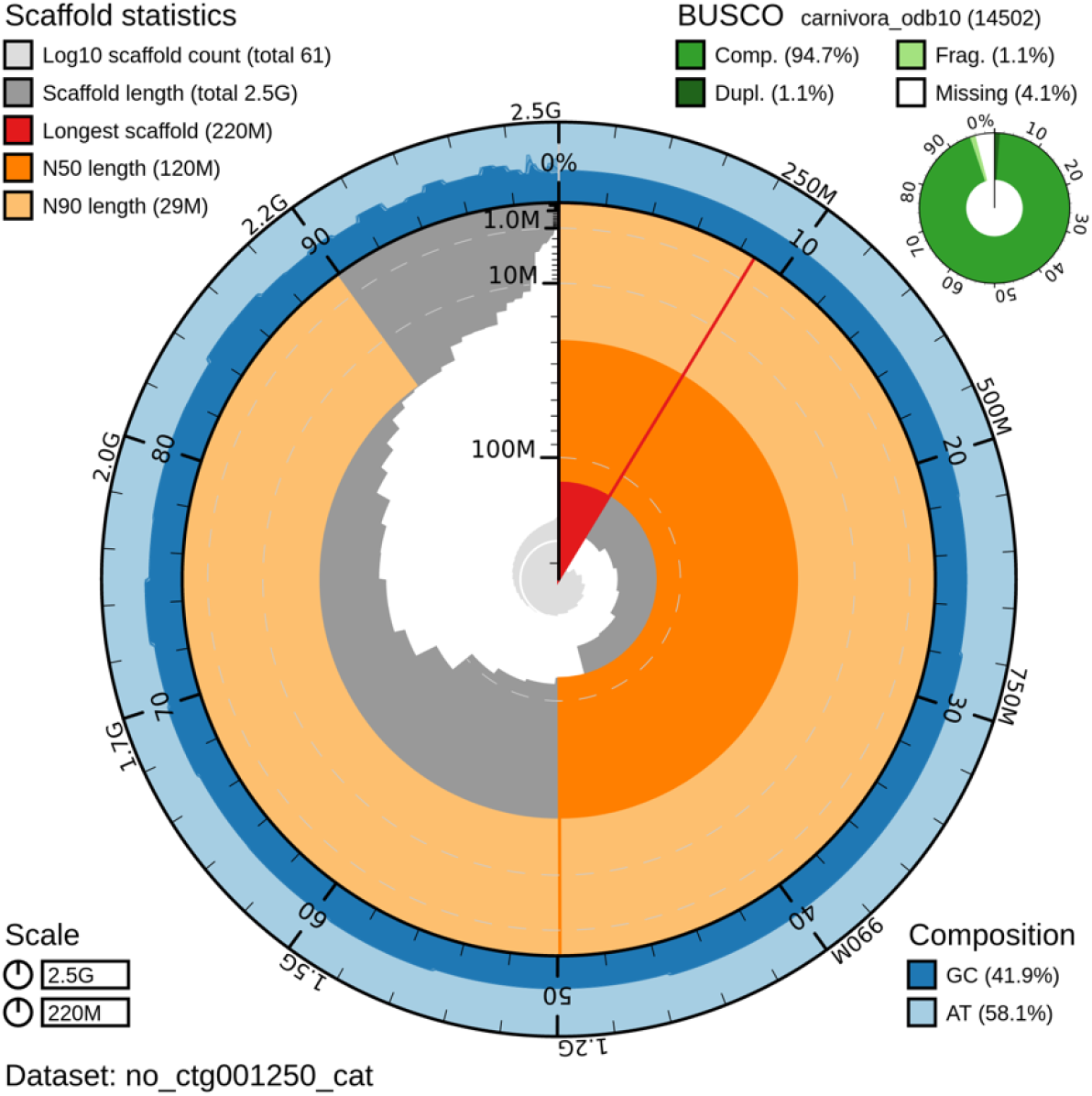
Snail plot of the contig-level *O. manul* nuclear genome assembly, OtoMan1.0. GC and AT content are shown in the outer ring and N50/N90 scores are shown in the inner ring. Results for benchmarking universal single-copy orthologs (BUSCO) genome quality scoring, including percentages of complete (Comp.), duplicated (Dupl.), fragmented (Frag.), and missing *Carnivora* genes, is included in the top right.

### DNA Methylation

Nanopore instruments can detect native DNA modifications via modified pore signals compared to the unmodified base. Here, 5-methylcytosine (5mCG) base modifications were called using Guppy with the contig-level assembly as an alignment reference, generating methylation data for 33,085,506 cytosine-guanine dinucleotides. Similar to other mammals^88^, global DNA methylation at CpG dinucleotides was high in *O. manul* at 78.2%.

Allele-specific methylation, a mechanism that facilitates allele-specific gene expression in genomic imprinting, was assessed by calling differentially methylated regions (DMRs) between *O. manul* pseudohaplotypes with DSS^54^. Strict calling parameters (i.e., methylation delta at least 50%, 100 bp length, at least 15 CpGs) yielded 91 unique loci (Table 4). Visualization of haplotagged read alignments at each DMR facilitated the discovery of 30 false positive loci, where apparent read misalignment for one of the two haplotypes led to a called DMR via a lack of CpGs on one allele. Of the original 91 loci, 61 (67%) were deemed true DMRs following manual inspection (Supplemental File 3). The validated DMRs had a mean length of 969 bp and a mean of 118 CpGs.

**Table 4:**
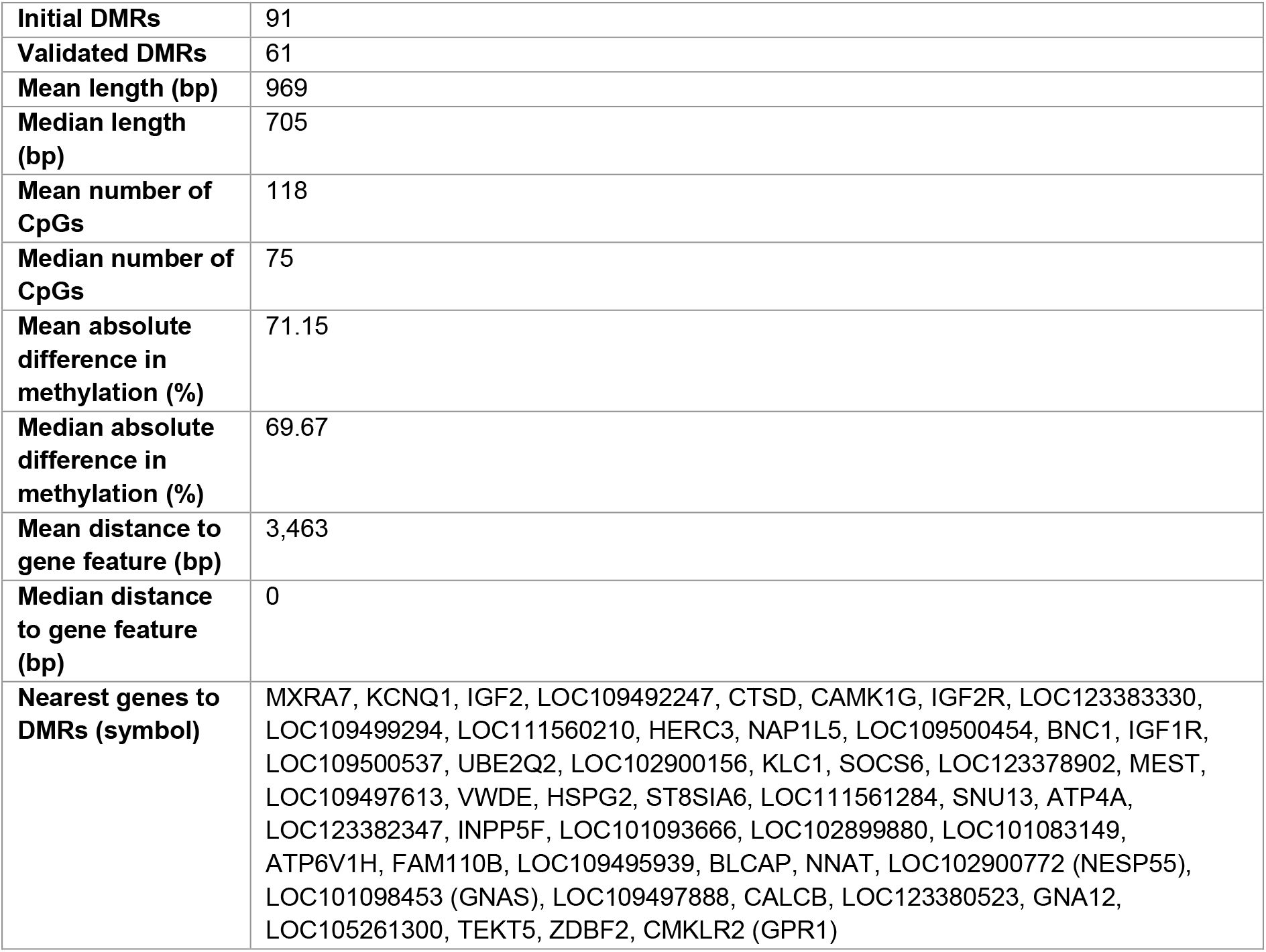
Summary statistics and nearest gene feature data for differentially methylated regions (DMRs) called between *O. manul* pseudohaplotyes. Gene annotation data were lifted onto the OtoMan1.0 assembly from *Felis catus* ([F.catus_Fca126_mat1.0](https://www.ncbi.nlm.nih.gov/data-hub/genome/GCF_018350175.1/)) and the nearest feature to each DMR was found with BEDTools^60^. Gene aliases are included in parenthesis.

Validated DMRs were annotated by lifting *F. catus* reference gene features onto the contig-level *O. manul* assembly and querying the nearest gene with BEDTools^60^. This method captured a number of classical imprinted loci described in humans and other animals, including GNAS, NNAT, IGF2, IGF2R, KCNQ1, MEST, and ZDBF2 (Table 4)^89–93^. In total, 41 validated DMRs directly overlapped 49 genes while non-overlapping DMRs fell between 165 bp and 56.7 kb away from the nearest feature. Putative novel imprinted loci, or DMRs near a feature not previously described as imprinted, including a 595 bp, 106-CpG DMR in the first exon of Von Willebrand factor D and EGF domain-containing protein (VWDE). Representative DMRs are shown in Figure 6. The mean absolute difference in methylation between the two pseudohaplotypes at DMRs was 71.15%. Most validated DMRs (44 of 61,72.1%) were hypomethylated on the alternate pseudohaplotype, while 17 were hypermethylated.

**Figure 6:**
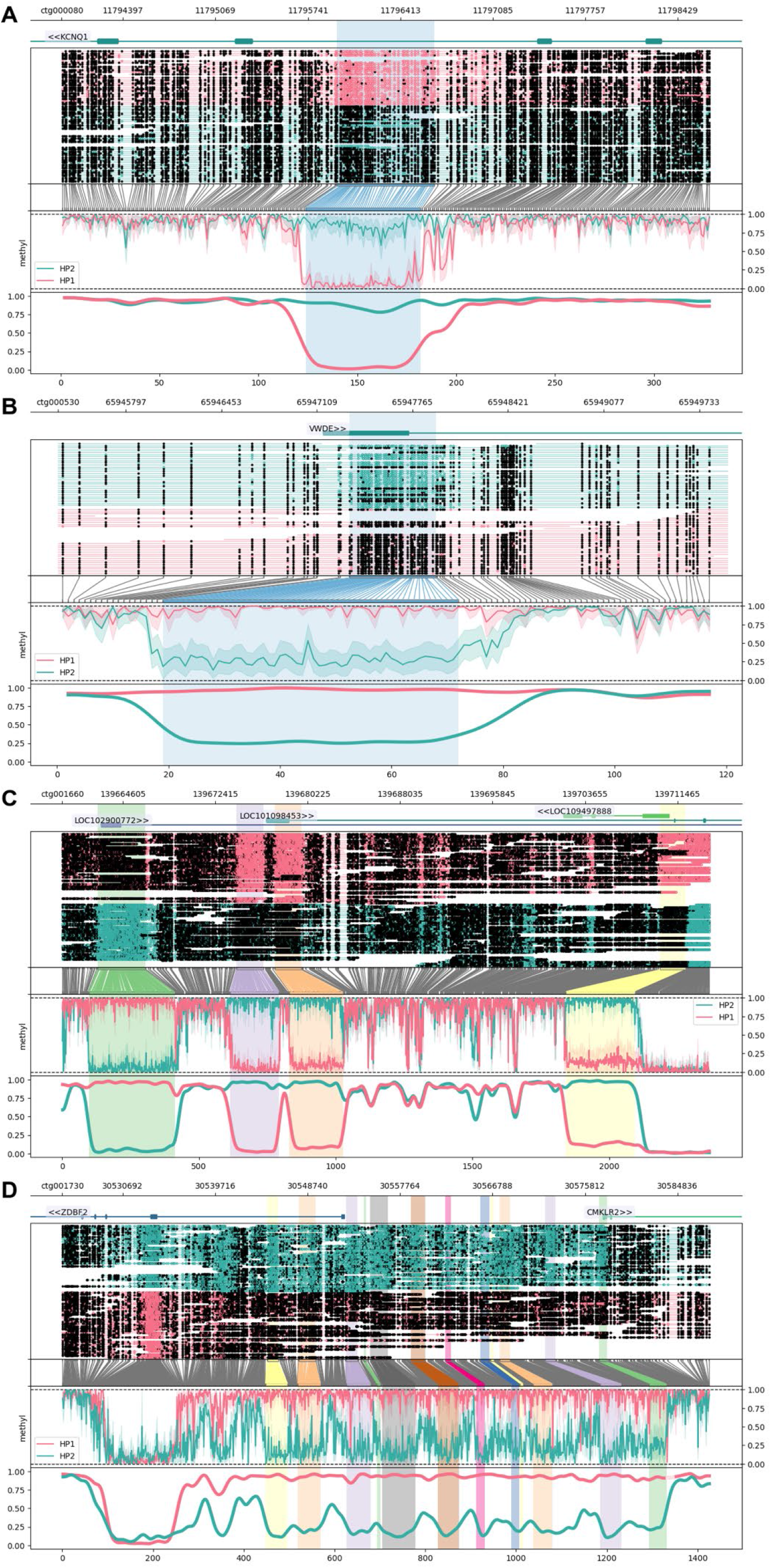
Allele-specific DNA methylation at differentially located loci (DMRs) called between *O. manul* assembly pseudohaplotypes assembled from a whole blood sample. DMRs were visualized with the package Methylartist^61^. Annotation of the 61 manually validated DMRs was accomplished by lifting gene coordinates from the *Felis catus* reference assembly ([F.catus_Fca126_mat1.0](https://www.ncbi.nlm.nih.gov/data-hub/genome/GCF_018350175.1/)) onto OtoMan1.0 and finding the nearest feature. The results revealed (A,B) singleton DMRs and (C,D) multi-DMR clusters within and near genes. Genes overlapping DMRs included previously described imprinted loci such as (A) potassium voltage-gated channel subfamily Q member 1 (KCNQ1) and (B) the guanine nucleotide-binding protein G(S) subunit alpha (GNAS) complex locus (aliases LOC101098453, LOC102900772). DMRs also overlapped genes not previously reported as imprinted, including (C) von Willebrand factor D and EGF domains (VWDE), a protein with predicted involvement in anatomical development^96^. (D) The most notable locus was a 33.2 kb region containing 12 DMRs all hypomethylated on pseudohaplotype 2. The DMRS overlapped the 5’ end of a classical imprinted gene, zinc finger DBF-type containing 2 (ZDBF2), on the negative strand and the 5’ end of chemerin chemokine-like receptor 2 (CMKLR2) on the positive strand. The CMKLR2 antisense RNA (CMKLR2AS; alias GPR1AS), which is not annotated in *F. catus*, exhibits imprinted expression in the human placenta^97^.

Of the 61 validated DMRs, 12 (19.67%) occurred in one contiguous 33.2 kb region on feline chromosome C1 (Figure 6). All DMRs in the region were hypomethylated on pseudohaplotype 2. The region fell directly upstream of 2 genes, chemerin chemokine-like receptor 2 (CMKLR2) on the positive strand and zinc finger DBF-type containing 2 (ZDBF2) on the negative strand. ZDBF2 is a canonical imprinted gene expressed paternally in most human and mouse tissues^94^; imprinted expression of CMKLR2, a paralog of opioid receptor kappa 1 (OPRK1), has not been reported, but expression of its antisense transcript (GPR1AS) is imprinted in human and mouse placenta^95^.

### Mitochondrial Genome

The mitochondrial genome assembly built using Flye in metagenome mode contained one circular contig 17,097 bp in length with 800x read coverage (Figure 7). Megablast revealed a 99.7% nucleotide identity match to a previously assembled *O. manul* mitogenome (Supplemental File 3)^98^. However, our assembly was longer than this previous manul mitogenome assembled from Sanger sequencing reads in 2019, which was 16,672 bp (GenBank MH978908.1). The 400 bp gap identified between our assembly and the 2019 version was located in the non-coding D-loop region of the mitogenome, making misassembly less likely than in the case of a large genic insertion. A 2015 Illumina mitogenome assembly for *O. manul* was closer to our assembly’s length at 17,009 bp (KR132585.1), but identity and query coverage were lower. From our mitogenome, a new phylogeny was built using tiger (*Panthera tigris*) as the outgroup. The resulting phylogeny matched previous nuclear DNA trees, with *Otocolobus* sharing its most recent ancestor with *Prionailurus*^14^ in contradiction of a 2016 mitochondrial phylogeny that placed *Otocolobus* closer to *Felis*^15^. The mitogenome was deposited as part of the scaffolded primary assembly at BioProject PRJNA885133.

**Figure 7:**
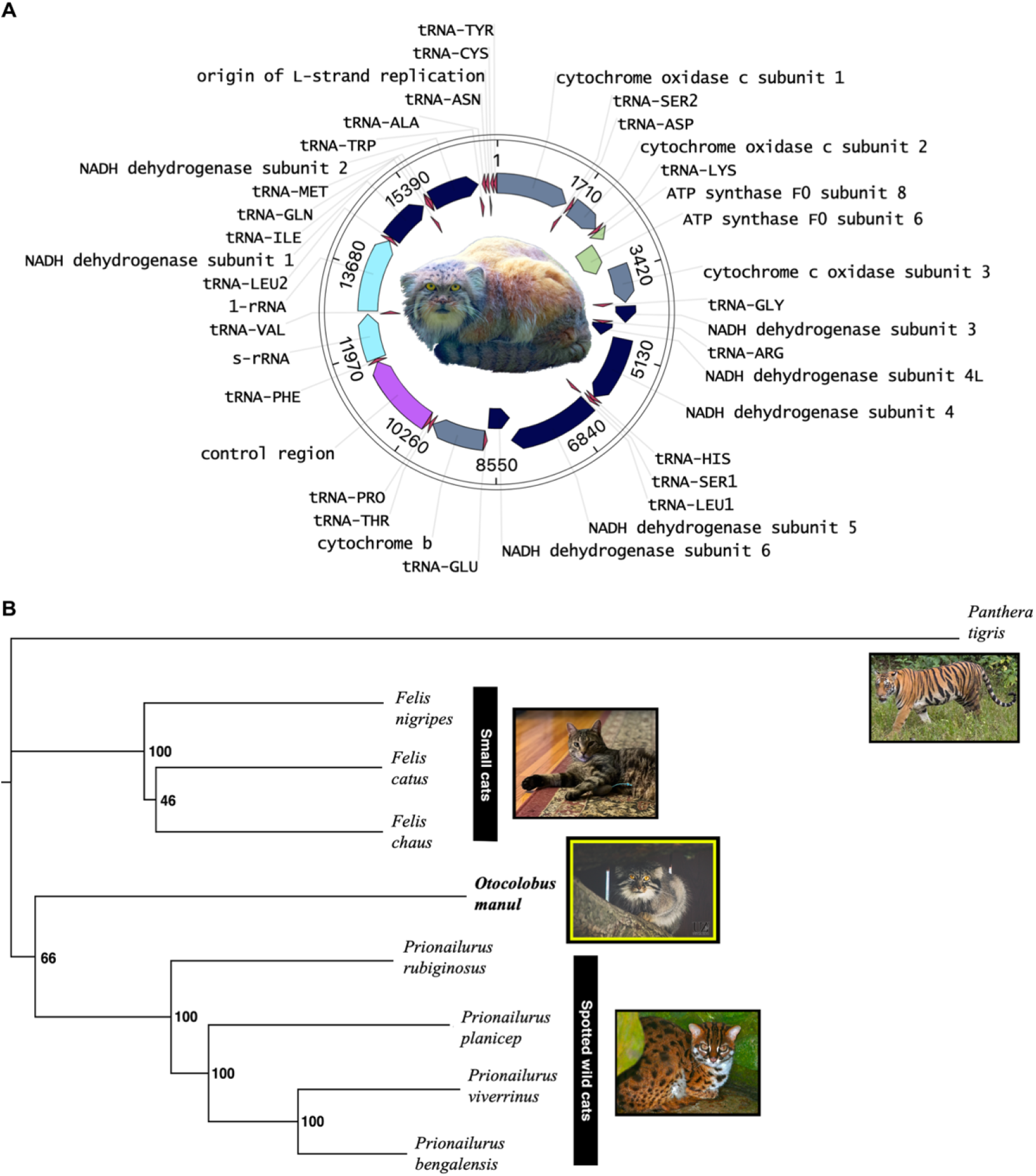
Nanopore mitochondrial genome assembly of *O. manul* and felid mtDNA phylogeny. (A) The circular length of the built *O. manul* mitogenome assembly was 17,097 bp; read coverage was 800x. (B) Phylogenetic relationships of *O. manul* were derived through bootstrapped maximum likelihood comparison of the assembled mitogenome to existing Felidae references. In contrast to previously published mtDNA for *O. manul*^99^, the current tree matches those generated through analysis of nuclear DNA^14^.

## DISCUSSION

Here, we have produced a highly contiguous diploid nuclear genome assembly, mitogenome, and allele-specific DNA methylation dataset for the manul cat (*Otocolobus manul*) via standalone nanopore sequencing. With 61 contigs and a contig N50 of 118.2 Mb, the assembly was more contiguous than any other *Felidae* reference genome to date before scaffolding. The felid clade is particularly well-suited for cross-genus scaffolding due to conserved genome collinearity among its members; our scaffolding strategy may not yield such unambiguous results in other animal families^80,83^. Comparing our diploid assembly to the original read set yielded a k-mer completeness score of 97.5% and quality value (QV) of 45.4, indicating a nucleotide error rate of less than 1 in 50,000 bases. This error rate is 5-fold better than the “platinum-quality” reference genome standards established by the Vertebrate Genomes Project (VGP), which suggest QV40, contig N50 ≥ 1 Mb, and chromosomal scaffold N50 ≥ 10 Mb^30,100^.

Our assembly’s BUSCO score for *CarniVora-*specific genes, 94.7%, also fell within 1% of the most complete felid genomes. Importantly, BUSCO values were re-calculated locally using an updated version of the program (v5.3.2), leading to lower scores overall compared to those currently displayed on NCBI’s Genome pages (v4.1.4). The described genome, OtoMan1.0, reflects the continued, rapid improvement of assembly pipelines and the necessity of long-read sequencing for generating contiguous, high-quality reference genomes^101–104^. Our results also provide evidence against concerns for low nanopore-only assembly quality due to sequencing error rates.

Repetitive elements have been minimally explored in felids despite their high genome occupancy and role in phenotype evolution^105–108^. SINE element insertions are major sources of genomic diversity in canids, with links to multiple phenotypes in domestic dogs (*Canis lupus familiaris*) including early retinal degeneration, polyneuropathy, myopathy, and merle coat color^109–115^. While global repeat content is strongly conserved across *Felidae* based on our analysis, individual repeat family activity is potentially variable and relevant to intra- and inter-species conservation. In a biomedically relevant example, domestic cats but not large cats possess an infectious endogenous retrovirus (ERV), RD-114, which can contaminate vaccines manufactured with feline cell lines and infect experimentally inoculated dogs^116–118^. Long-read sequencing enables more accurate investigation of genomic repeat content; single reads can span full TEs and capture unique flanking host sequences, reducing multi-mapping issues inherent in short read assembly^119–121^.

Nanopore sequencing permits native DNA methylation detection alongside DNA base calls, combining genome assembly and epigenome analysis into one step at the bench top. The epigenome is responsive to an animal’s environment and shifts in a predictable manner across the life span^122,123^. Here, we present a phased methylome and allele-specific DNA methylation analysis for *O. manul*. Our differentially methylated region (DMR) detection criteria were relatively strict in an effort to reduce false positives and yielded 61 validated loci. We successfully captured a number of classical imprinted genes including NESP55/GNAS, KCNQ1, NNAT, IGF1R, and IGF2/H19^124–128^. IGF2R contained a DMR in this study suggestive of imprinted expression; this gene is maternally expressed in dogs^129,130^ while monoallelic expression appears to have been lost in the primate lineage^131^. Putative novel imprinting loci were also identified in genes such as Von Willebrand factor D and EGF domain-containing protein (VWDE), a paralog of fibulin 2 (FBLN2) required for multi-tissue limb regeneration in the axolotl (*Ambystoma mexicanum)^132^*. A 2012 study found that 40% of human genomes deposited by the 1000 Genomes project (www.1000genomes.org) carried at least one loss-of-function mutation in VWDE^133^. This locus and others were evaluated as distinct from 30 false DMRs, where misassembly resulted in a lack of CpGs on one pseudohaplotype. This phenomenon is not detectable to the DMR calling software *DSS*, necessitating manual curation; humans have over 200 imprinted genes^127,134^ meaning our DMR calling parameters likely also introduced false negatives. Although imprinted expression cannot be predicted based on allele-specific methylation alone, expanded investigations of this phenomenon in diverse, non-domesticated species via nanopore sequencing will enrich our understanding of the mechanism.

Detection of low genome heterozygosity in this captive-bred *O. manul* individual is a concerning but valuable insight for both zoo breeding programs and wild population management. At 0.048%, our estimate of heterozygosity using biallelic pseudohaplotype SNVs was similar to the critically endangered Amur tiger (*Panthera tigris altaica*), a species that experienced a severe population bottleneck in the 1940s with a current estimated population of 400 animals^135–138^. In contrast, the manul cat population is estimated at 58,000 individuals^1^. Although captive breeding programs are a crucial tool in global conservation, a small pool of founder animals inevitably leads to reduced genetic diversity. Practices like sperm biobanking have the potential to support captive population genetic quality, but such systems are underutilized^139^. Fortunately, our estimate of effective population size using runs of homozygosity was relatively large at N_e_ = 10,000. Results also suggested that manul cats experienced a crash in genetic diversity, but that this event was in the relatively distant past (~10,000 years ago); effective population size appears to have continued recovering despite anthropogenic challenges to the species. Given the large, low-density geographic range and solitary social strategy of *O. manul*, it is unclear whether these values are typical across the entire population.

While largely conserved among metazoans, mitogenomes also contain structural and nucleotide variation that can serve as phylogenetic markers^140^. Their smaller size, relative to nuclear genomes, make them an attractive tool in phylogenetics. However, incongruities between mitogenome and nuclear DNA phylogenies can occur due to uniparental inheritance of organelles^141^, horizontal gene transfer^142^, or incomplete lineage sorting^143^. Previously, an *O. manul* mitogenome assembly placed *Otocolobus* as a sister taxa to the *Felis* clade^15^. The mitogenome assembly presented here contains a previously unidentified 400 bp gap in the D-loop control region; a new phylogenetic analysis placed *O. manul* closest to *Prionailurus* in agreement with the nuclear genome phylogeny^14^. This finding suggests that *O. manul* nuclear DNA and mtDNA have similar phylogenetic histories. Recently, felid mitogenomes were used to investigate the contribution of a wild cat to domestication of *F. catus^144^*. Using our mitogenome assembly, a similar study could investigate introgression among wild Asian cats (i.e., *O. manul* and *Prionailurus spp.*) and between wild and domestic cats in this region. Hybridization among cat species could have important implications for genetic diversity and conservation. Notably, evidence of hybridization among domestic and wild cats is found in South Africa and Europe, and is related to human population density in proximate spaces to wild cat habitat^145,146^.

Long-read sequencing in general has been critical for increased genome assembly quality, but the portability and cost-effectiveness of nanopore sequencing make it particularly valuable for reducing barriers to entry in genomics^82,104^. Still, assembly of mammalian-sized genomes (3+ Gb) can exceed computational and financial resources available to small laboratories. Even Flye, an efficient assembler^31^, required distributed computing resources when using our full read set (Table 2; Supplemental File 3). Unequal access to such resources can inadvertently perpetuate “parachute science” and limit the participation of researchers from historically colonized countries in endemic species conservation^147–149^. However, we were able to build *O. manul* assemblies locally using a subset of the longest reads with only a small reduction in BUSCO score (Supplemental File 3). On-demand cloud computing resources such as Amazon Web Services (AWS) are also increasingly affordable, eliminating the necessity of access to a university cluster. We hope that the provided data and computational pipeline will be useful resources for *O. manul* conservation as well as for other small groups assembling mammal genomes.

## DATA AVAILABILITY

DNA sequencing reads are available in FASTQ format in the Sequencing Read Archive repository (https://www.ncbi.nlm.nih.gov/sra) under submission SRR22085263. The scaffolded primary assembly (including mitogenome) is available as BioProject PRJNA885133; the contig-level alternate assembly is available as BioProject PRJNA889808. Modified base calls (DNA methylation) are also available under BioProject PRJNA885133. Sample data are available as BioSample SAMN31076064.

## ACKNOWLEDGEMENTS

We thank the Utica Zoo and Tater for providing the whole blood sample used in this study. We also thank Dr. Adam Ewing for assistance with the program Methylartist. For Figure 7B, leopard cat photo credit goes to [Bernard Dupont](https://creativecommons.org/licenses/by-sa/2.0/) and tiger photo credit to Charles J. Sharp, both under Creative Commons licenses.

## REFERENCES

1. Otocolobus manul: Ross, s., Barashkova, a., Dhendup, t., Munkhtsog, b., Smelansky, i., Barclay, d. & moqanaki, e. (2019) doi:10.2305/iucn.uk.2020-2.rlts.t15640a180145377.en.

2. Gittleman, J. L. Heptner, V. G., and Sludskii, A. A. 1992. MAMMALS OF THE SOVIET UNION. VOLUME II, PART 2. CARNIVORA (HYAENAS AND CATS). Smithsonian Institution Libraries and National Science Foundation. Journal of Mammalogy 74, 510–511 (1993).

3. Murdoch, J., Tserendorj, M. & Reading, R. Pallas’ cat ecology and conservation in the semi-desert steppes of mongolia. CAT News 45, 18–19 (2006).

4. BBC. The grumpiest cat in the world. (2022). https://www.bbc.co.uk/programmes/p0cz5p0y. Accessed November 30, 2022.

5. Ross, S., Munkhtsog, B. & Harris, S. Dietary composition, plasticity, and prey selection of Pallas’s cats. Journal of Mammalogy 91, 811–817 (2010).

6. Pallas, P. S. Reise durch verschiedene provinzen des russischen reichs. Vol. Reise aus sibirien zurück an die wolga im 1773ten jahr. in (St. Petersburg, Russian Empire: Kayserliche Academie der Wissenschafen, 1776).

7. Brandt, J. Observations sur le manoul (felis manul pallas). Bull. sc. Ac. Imp. Sc. St. Petersb 9, 37–39 (1842).

8. Pocock, R. I. Exhibition of a photograph and the skull of a specimen of the manul or pallas’ cat (felis manul) that had recently died in the society’s menagerie with some remarks on the species. Proceedings of the Zoological Society of London 299–306 (1907).

9. Otocolobus manul: Ross, s., Barashkova, a., Farhadinia, m.s., Appel, a., Riordan, p., Sanderson, j. & munkhtsog, b. (2014) doi:10.2305/iucn.uk.2016-1.rlts.t15640a87840229.en.

10. Spong, G., Johansson, M. & Björklund, M. High genetic variation in leopards indicates large and long-term stable effective population size. Molecular Ecology 9, 1773–1782 (2000).

11. Palstra, F. P. & Ruzzante, D. E. Genetic estimates of contemporary effective population size: what can they tell us about the importance of genetic stochasticity for wild population persistence? Molecular Ecology 17, 3428–3447 (2008).

12. Ross, S., Munkhtsog, B. & Harris, S. Dietary composition, plasticity, and prey selection of Pallas’s cats. Journal of Mammalogy 91, 811–817 (2010).

13. Barclay, D., Smelansky, I., Nygren, E. & Antonevich, A. Legal status, utilisation, management and conservation of manul. Cat News Special 37–40 (2019).

14. Johnson, W. E. et al. The Late Miocene Radiation of Modern Felidae: A Genetic Assessment. Science 311, 73–77 (2006).

15. Li, G., Davis, B. W., Eizirik, E. & Murphy, W. J. Phylogenomic evidence for ancient hybridization in the genomes of living cats (Felidae). Genome Research 26, 1–11 (2015).

16. Sakamoto, M. & Ruta, M. Convergence and Divergence in the Evolution of Cat Skulls: Temporal and Spatial Patterns of Morphological Diversity. PLoS ONE 7, e39752 (2012).

17. Mohamed, M. et al. A Transposon Story: From TE Content to TE Dynamic Invasion of Drosophila Genomes Using the Single-Molecule Sequencing Technology from Oxford Nanopore. Cells 9, 1776 (2020).

18. Moss, E. L., Maghini, D. G. & Bhatt, A. S. Complete, closed bacterial genomes from microbiomes using nanopore sequencing. Nature Biotechnology 38, 701–707 (2020).

19. Ewing, A. D. et al. Nanopore Sequencing Enables Comprehensive Transposable Element Epigenomic Profiling. Molecular Cell 80, 915–928.e5 (2020).

20. Razin, A. & Cedar, H. DNA methylation and gene expression. Microbiological Reviews 55, 451–458 (1991).

21. Tate, P. H. & Bird, A. P. Effects of DNA methylation on DNA-binding proteins and gene expression. Current Opinion in Genetics & Development 3, 226–231 (1993).

22. Greenberg, M. V. C. & Bourc’his, D. The diverse roles of DNA methylation in mammalian development and disease. Nature Reviews Molecular Cell Biology 20, 590–607 (2019).

23. Zemach, A., McDaniel, I. E., Silva, P. & Zilberman, D. Genome-Wide Evolutionary Analysis of Eukaryotic DNA Methylation. Science 328, 916–919 (2010).

24. Doskočil, J. & Šorm, F. Distribution of 5-methylcytosine in pyrimidine sequences of deoxyribonucleic acids. Biochimica et Biophysica Acta 55, 953–959 (1962).

25. Riggs, A. D. X inactivation, differentiation, and DNA methylation. Cytogenetic and Genome Research 14, 9–25 (1975).

26. Cooper, D. N. & Krawczak, M. Cytosine methylation and the fate of CpG dinucleotides in vertebrate genomes. Human Genetics 83, 181–188 (1989).

27. Shafin, K. et al. Haplotype-aware variant calling with PEPPER-Margin-DeepVariant enables high accuracy in nanopore long-reads. Nature Methods 18, 1322–1332 (2021).

28. Battaglia, S. et al. Long-range phasing of dynamic, tissue-specific and allele-specific regulatory elements. Nature Genetics 54, 1504–1513 (2022).

29. Akbari, V. et al. Genome-wide detection of imprinted differentially methylated regions using nanopore sequencing. eLife 11, (2022).

30. Rhie, A. et al. Towards complete and error-free genome assemblies of all vertebrate species. (2020).

31. Kolmogorov, M., Yuan, J., Lin, Y. & Pevzner, P. A. Assembly of long, error-prone reads using repeat graphs. Nature Biotechnology 37, 540–546 (2019).

32. Shafin, K. et al. Nanopore sequencing and the Shasta toolkit enable efficient de novo assembly of eleven human genomes. Nature Biotechnology 38, 1044–1053 (2020).

33. Vaser, R. & Šikić, M. Time-and memory-efficient genome assembly with Raven. Nature Computational Science 1, 332–336 (2021).

34. Vaser, R., Sović, I., Nagarajan, N. & Šikić, M. Fast and accurate de novo genome assembly from long uncorrected reads. Genome Research 27, 737–746 (2017).

35. Hu, J., Fan, J., Sun, Z. & Liu, S. NextPolish: a fast and efficient genome polishing tool for long-read assembly. Bioinformatics 36, 2253–2255 (2019).

36. Manni, M., Berkeley, M. R., Seppey, M. & Zdobnov, E. M. BUSCO: Assessing Genomic Data Quality and Beyond. Current Protocols 1, (2021).

37. Manni, M., Berkeley, M. R., Seppey, M., Simão, F. A. & Zdobnov, E. M. BUSCO Update: Novel and Streamlined Workflows along with Broader and Deeper Phylogenetic Coverage for Scoring of Eukaryotic, Prokaryotic, and Viral Genomes. Molecular Biology and Evolution 38, 4647–4654 (2021).

38. Solares, E. A. et al. Rapid Low-Cost Assembly of the *Drosophila melanogaster* Reference Genome Using Low-Coverage, Long-Read Sequencing. G3 Genes|Genomes|Genetics 8, 3143–3154 (2018).

39. Solares, E. A. et al. Rapid Low-Cost Assembly of the *Drosophila melanogaster* Reference Genome Using Low-Coverage, Long-Read Sequencing. G3 Genes|Genomes|Genetics 8, 3143–3154 (2018).

40. Roach, M. J., Schmidt, S. A. & Borneman, A. R. Purge Haplotigs: allelic contig reassignment for third-gen diploid genome assemblies. BMC Bioinformatics 19, (2018).

41. Wood, D. E., Lu, J. & Langmead, B. Improved metagenomic analysis with Kraken 2. Genome Biology 20, (2019).

42. Breitwieser, F. P. & Salzberg, S. L. Pavian: interactive analysis of metagenomics data for microbiome studies and pathogen identification. Bioinformatics 36, 1303–1304 (2019).

43. Zhang, Z., Schwartz, S., Wagner, L. & Miller, W. A Greedy Algorithm for Aligning DNA Sequences. Journal of Computational Biology 7, 203–214 (2000).

44. Challis, R., Richards, E., Rajan, J., Cochrane, G. & Blaxter, M. BlobToolKit Interactive Quality Assessment of Genome Assemblies. G3 Genes|Genomes|Genetics 10, 1361–1374 (2020).

45. Pedersen, B. S. & Quinlan, A. R. Mosdepth: quick coverage calculation for genomes and exomes. Bioinformatics 34, 867–868 (2017).

46. Rhie, A., Walenz, B. P., Koren, S. & Phillippy, A. M. Merqury: reference-free quality, completeness, and phasing assessment for genome assemblies. Genome Biology 21, (2020).

47. Storer, J., Hubley, R., Rosen, J., Wheeler, T. J. & Smit, A. F. The Dfam community resource of transposable element families, sequence models, and genome annotations. Mobile DNA 12, (2021).

48. Keilwagen, J. et al. Using intron position conservation for homology-based gene prediction. Nucleic Acids Research 44, e89–e89 (2016).

49. Keilwagen, J., Hartung, F., Paulini, M., Twardziok, S. O. & Grau, J. Combining RNA-seq data and homology-based gene prediction for plants, animals and fungi. BMC Bioinformatics 19, (2018).

50. Martin, M. et al. WhatsHap: Fast and accurate read-based phasing. (2016).

51. Danecek, P. et al. The variant call format and VCFtools. Bioinformatics 27, 2156–2158 (2011).

52. Danecek, P. et al. Twelve years of SAMtools and BCFtools. GigaScience 10, (2021).

53. Terhorst, J., Kamm, J. A. & Song, Y. S. Robust and scalable inference of population history from hundreds of unphased whole genomes. Nature Genetics 49, 303–309 (2016).

54. Wu, H., Wang, C. & Wu, Z. A new shrinkage estimator for dispersion improves differential expression detection in RNA-seq data. Biostatistics 14, 232–243 (2012).

55. Feng, H., Conneely, K. N. & Wu, H. A Bayesian hierarchical model to detect differentially methylated loci from single nucleotide resolution sequencing data. Nucleic Acids Research 42, e69–e69 (2014).

56. Wu, H. et al. Detection of differentially methylated regions from whole-genome bisulfite sequencing data without replicates. Nucleic Acids Research gkv715 (2015) doi:10.1093/nar/gkv715.

57. Park, Y. & Wu, H. Differential methylation analysis for BS-seq data under general experimental design. Bioinformatics 32, 1446–1453 (2016).

58. Shumate, A. & Salzberg, S. L. Liftoff: accurate mapping of gene annotations. Bioinformatics 37, 1639–1643 (2021).

59. Quinlan, A. R. & Hall, I. M. BEDTools: a flexible suite of utilities for comparing genomic features. Bioinformatics 26, 841–842 (2010).

60. Quinlan, A. R. BEDTools: The Swiss-Army Tool for Genome Feature Analysis. Current Protocols in Bioinformatics 47, (2014).

61. Cheetham, S. W., Kindlova, M. & Ewing, A. D. Methylartist: tools for visualizing modified bases from nanopore sequence data. Bioinformatics 38, 3109–3112 (2022).

62. Lin, J.-H., Chen, L.-C., Yu, S.-C. & Huang, Y.-T. LongPhase: an ultra-fast chromosome-scale phasing algorithm for small and large variants. Bioinformatics 38, 1816–1822 (2022).

63. Diesh, C. et al. JBrowse 2: A modular genome browser with views of synteny and structural variation. (2022).

64. Lopez, J. V., Cevario, S. & O’Brien, S. J. Complete Nucleotide Sequences of the Domestic Cat (Felis catus) Mitochondrial Genome and a Transposed mtDNA Tandem Repeat (Numt) in the Nuclear Genome. Genomics 33, 229–246 (1996).

65. Li, H. Minimap2: pairwise alignment for nucleotide sequences. Bioinformatics 34, 3094–3100 (2018).

66. Wanner, N., Larsen, P. A., McLain, A. & Faulk, C. The mitochondrial genome and Epigenome of the Golden lion Tamarin from fecal DNA using Nanopore adaptive sequencing. BMC Genomics 22, (2021).

67. Edgar, R. C. MUSCLE: multiple sequence alignment with high accuracy and high throughput. Nucleic Acids Research 32, 1792–1797 (2004).

68. Nguyen, L.-T., Schmidt, H. A., Haeseler, A. von & Minh, B. Q. IQ-TREE: A Fast and Effective Stochastic Algorithm for Estimating Maximum-Likelihood Phylogenies. Molecular Biology and Evolution 32, 268–274 (2014).

69. Alonge, M. et al. Automated assembly scaffolding elevates a new tomato system for high-throughput genome editing. (2021).

70. Marçais, G. et al. MUMmer4: A fast and versatile genome alignment system. PLOS Computational Biology 14, e1005944 (2018).

71. Pontius, J. U. etal. Initial sequence and comparative analysis of the cat genome. Genome Research 17, 1675–1689 (2007).

72. Buckley, R. M. et al. A new domestic cat genome assembly based on long sequence reads empowers feline genomic medicine and identifies a novel gene for dwarfism. PLOS Genetics 16, e1008926 (2020).

73. Brashear, W. A., Bredemeyer, K. R. & Murphy, W. J. Genomic architecture constrained placental mammal X Chromosome evolution. Genome Research 31, 1353–1365 (2021).

74. Burger, P. A. et al. Analysis of the mitochondrial genome of cheetahs (Acinonyx jubatus) with neurodegenerative disease. Gene 338, 111–119 (2004).

75. Prost, S. et al. Genomic analyses show extremely perilous conservation status of African and Asiatic cheetahs (*Acinonyx jubatus*). Molecular Ecology 31, 4208–4223 (2022).

76. Bredemeyer, K. R. et al. Rapid Macrosatellite Evolution Promotes X-Linked Hybrid Male Sterility in a Feline Interspecies Cross. Molecular Biology and Evolution 38, 5588–5609 (2021).

77. Rhie, A. et al. Towards complete and error-free genome assemblies of all vertebrate species. Nature 592, 737–746 (2021).

78. Abascal, F. et al. Extreme genomic erosion after recurrent demographic bottlenecks in the highly endangered Iberian lynx. Genome Biology 17, (2016).

79. Lei, W., XiaoBing, W., Zhu, L. & Jiang, Z. Mitogenomic analysis of the genus Panthera. Science China Life Sciences 54, 917–930 (2011).

80. Bredemeyer, K. R. et al. Ultracontinuous Single Haplotype Genome Assemblies for the *Domestic Cat (*Felis catus*) and Asian Leopard Cat (Prionailurus bengalensis*). Journal of Heredity 112, 165–173 (2020).

81. Tamazian, G. et al. *Draft *de novo* Genome Assembly of the Elusive Jaguarundi, Puma yagouaroundi*. Journal of Heredity 112, 540–548 (2021).

82. Faulk, C. *De novo* sequencing, diploid assembly, and annotation of the black carpenter ant, *Camponotus pennsylvanicus*, and its symbionts by one person for $1000, using nanopore sequencing. Nucleic Acids Research (2022) doi:10.1093/nar/gkac510.

83. Armstrong, E. E. et al. Long live the king: chromosome-level assembly of the lion (Panthera leo) using linked-read, Hi-C, and long-read data. BMC Biology 18, (2020).

84. Schrader, L. & Schmitz, J. The impact of transposable elements in adaptive evolution. Molecular Ecology 28, 1537–1549 (2018).

85. Böhne, A., Brunet, F., Galiana-Arnoux, D., Schultheis, C. & Volff, J.-N. Transposable elements as drivers of genomic and biological diversity in vertebrates. Chromosome Research 16, 203–215 (2008).

86. Platt, R. N., Vandewege, M. W. & Ray, D. A. Mammalian transposable elements and their impacts on genome evolution. Chromosome Research 26, 25–43 (2018).

87. Meredith, R. W. et al. Impacts of the Cretaceous Terrestrial Revolution and KPg Extinction on Mammal Diversification. Science 334, 521–524 (2011).

88. Ehrlich, M. et al. Amount and distribution of 5-methylcytosine in human DNA from different types of tissues or cells. Nucleic Acids Research 10, 2709–2721 (1982).

89. Bastepe, M. et al. Deletion of the NESP55 differentially methylated region causes loss of maternal GNAS imprints and pseudohypoparathyroidism type Ib. Nature Genetics 37, 25–27 (2004).

90. Zaitoun, I. & Khatib, H. Assessment of genomic imprinting of SLC38A4, NNAT, NAP1L5, and H19 in cattle. BMC Genetics 7, (2006).

91. Brabazon, D. C., Callanan, J. J. & Nolan, C. M. Imprinting of canine *IGF2* and *H19*. Animal Genetics 53, 108–118 (2021).

92. Eßinger, C. et al. Frequency of KCNQ1 variants causing loss of methylation of Imprinting Centre 2 in Beckwith-Wiedemann syndrome. Clinical Epigenetics 12, (2020).

93. Li, X. et al. Isoform-specific imprinting of the MEST gene in porcine parthenogenetic fetuses. Gene 558, 287–290 (2015).

94. Kobayashi, H. et al. Identification of the mouse paternally expressed imprinted gene Zdbf2 on chromosome 1 and its imprinted human homolog ZDBF2 on chromosome 2. Genomics 93, 461–472 (2009).

95. Kobayashi, H. et al. Epigenetic and transcriptional features of the novel human imprinted *lncRNA*GPR1AS*suggest it is a functional ortholog to mouseZdbf2linc*. Epigenetics 8, 635–645 (2013).

96. Iwata, K. et al. von Willebrand factor D and EGF domains regulate ameloblast differentiation and enamel formation. Journal of Cellular Physiology 237, 1964–1979 (2021).

97. Kobayashi, H. et al. Epigenetic and transcriptional features of the novel human imprinted *lncRNA*GPR1AS*suggest it is a functional ortholog to mouseZdbf2linc*. Epigenetics 8, 635–645 (2013).

98. Xu, Y. et al. The complete mitochondrial genome of Pallas’s cat (Otocolobus manul). Mitochondrial DNA Part B 4, 658–659 (2019).

99. Xu, Y. et al. The complete mitochondrial genome of Pallas’s cat (Otocolobus manul). Mitochondrial DNA Part B 4, 658–659 (2019).

100. Morin, P. A. et al. Reference genome and demographic history of the most endangered marine mammal, the vaquita. Molecular Ecology Resources 21, 1008–1020 (2020).

101. Murigneux, V. et al. Comparison of long-read methods for sequencing and assembly of a plant genome. GigaScience 9, (2020).

102. Wick, R. R. & Holt, K. E. Benchmarking of long-read assemblers for prokaryote whole genome sequencing. F1000Research 8, 2138 (2019).

103. Chen, Z., Erickson, D. L. & Meng, J. Benchmarking Long-Read Assemblers for Genomic Analyses of Bacterial Pathogens Using Oxford Nanopore Sequencing. International Journal of Molecular Sciences 21, 9161 (2020).

104. Hotaling, S., Kelley, J. L. & Frandsen, P. B. Toward a genome sequence for every animal: Where are we now? Proceedings of the National Academy of Sciences 118, (2021).

105. Bhat, A. et al. Role of Transposable Elements in Genome Stability: Implications for Health and Disease. International Journal of Molecular Sciences 23, 7802 (2022).

106. Lavialle, C. et al. Paleovirology of ‘ *syncytins* retroviral *env* genes exapted for a role in placentation. Philosophical Transactions of the Royal Society B: Biological Sciences 368, 20120507 (2013).

107. Stoye, J. P. Studies of endogenous retroviruses reveal a continuing evolutionary saga. Nature Reviews Microbiology 10, 395–406 (2012).

108. Chiu, E. S. & VandeWoude, S. Presence of Endogenous Viral Elements Negatively Correlates with Feline Leukemia Virus Susceptibility in Puma and Domestic Cat Cells. Journal of Virology 94, (2020).

109. Walters-Conte, K. B., Johnson, D. L. E., Allard, M. W. & Pecon-Slattery, J. Carnivore-Specific SINEs (Can-SINEs): Distribution, Evolution, and Genomic Impact. Journal of Heredity 102, S2–S10 (2011).

110. Wiedmer, M. et al. A *RAB3GAP1* SINE Insertion in Alaskan Huskies with Polyneuropathy, Ocular Abnormalities, and Neuronal Vacuolation (POANV) Resembling Human Warburg Micro Syndrome 1 (WARBM1). G3 Genes|Genomes|Genetics 6, 255–262 (2016).

111. Goldstein, O., Kukekova, A. V., Aguirre, G. D. & Acland, G. M. Exonic SINE insertion in STK38L causes canine early retinal degeneration (erd). Genomics 96, 362–368 (2010).

112. Pelé, M., Tiret, L., Kessler, J.-L., Blot, S. & Panthier, J.-J. SINE exonic insertion in the PTPLA gene leads to multiple splicing defects and segregates with the autosomal recessive centronuclear myopathy in dogs. Human Molecular Genetics 14, 1417–1427 (2005).

113. Wang, W. & Kirkness, E. F. Short interspersed elements (SINEs) are a major source of canine genomic diversity. Genome Research 15, 1798–1808 (2005).

114. Clark, L. A., Wahl, J. M., Rees, C. A. & Murphy, K. E. Retrotransposon insertion in *SILV* is responsible for merle patterning of the domestic dog. Proceedings of the National Academy of Sciences 103, 1376–1381 (2006).

115. Murphy, S. C., Evans, J. M., Tsai, K. L. & Clark, L. A. Length variations within the Merle retrotransposon of canine PMEL: correlating genotype with phenotype. Mobile DNA 9, (2018).

116. Okada, M., Yoshikawa, R., Shojima, T., Baba, K. & Miyazawa, T. Susceptibility and production of a feline endogenous retrovirus (RD-114 virus) in various feline cell lines. Virus Research 155, 268–273 (2011).

117. Okabe, H., Gilden, R. V. & Hatanaka, M. RD 114 Virus-Specific Sequences in Feline Cellular RNA: Detection and Characterization. Journal of Virology 12, 984–994 (1973).

118. Yoshikawa, R., Shimode, S., Sakaguchi, S. & Miyazawa, T. Contamination of live attenuated vaccines with an infectious feline endogenous retrovirus (RD-114 virus). Archives of Virology 159, 399–404 (2013).

119. Yasir, M. et al. Long-read sequencing for identification of insertion sites in large transposon mutant libraries. Scientific Reports 12, (2022).

120. Logsdon, G. A., Vollger, M. R. & Eichler, E. E. Long-read human genome sequencing and its applications. Nature Reviews Genetics 21, 597–614 (2020).

121. Miga, K. H. et al. Telomere-to-telomere assembly of a complete human X chromosome. Nature 585, 79–84 (2020).

122. Raj, K. et al. Epigenetic clock and methylation studies in cats. GeroScience 43, 2363–2378 (2021).

123. Horvath, S. DNA methylation age of human tissues and cell types. Genome Biology 14, R115 (2013).

124. Ekström, T. J. Parental Imprinting and the *IGF2* Gene. Hormone Research 42, 176–181 (1994).

125. Tucci, V. et al. Genomic Imprinting and Physiological Processes in Mammals. Cell 176, 952–965 (2019).

126. Bartolomei, M. S., Zemel, S. & Tilghman, S. M. Parental imprinting of the mouse H19 gene. Nature 351, 153–155 (1991).

127. Jima, D. D. et al. Genomic map of candidate human imprint control regions: the imprintome. Epigenetics 17, 1920–1943 (2022).

128. Kanduri, C. et al. A Differentially Methylated Imprinting Control Region within the Kcnq1 Locus Harbors a Methylation-sensitive Chromatin Insulator. Journal of Biological Chemistry 277, 18106–18110 (2002).

129. O’Sullivan, F. M. et al. Imprinted expression of the canine IGF2R, in the absence of an anti-sense transcript or promoter methylation. Evolution & Development 9, 579–589 (2007).

130. Nolan, C., O’Sullivan, F., Brabazon, D. & Callanan, J. Genomic Imprinting inCanis familiaris. Reproduction in Domestic Animals 44, 16–21 (2009).

131. Killian, J. K. Divergent evolution in M6P/IGF2R imprinting from the jurassic to the quaternary. Human Molecular Genetics 10, 1721–1728 (2001).

132. Leigh, N. D. et al. von Willebrand factor D and EGF domains is an evolutionarily conserved and required feature of blastemas capable of multitissue appendage regeneration. Evolution & Development 22, 297–311 (2020).

133. MacArthur, D. G. et al. A Systematic Survey of Loss-of-Function Variants in Human Protein-Coding Genes. Science 335, 823–828 (2012).

134. Tucci, V. et al. Genomic Imprinting and Physiological Processes in Mammals. Cell 176, 952–965 (2019).

135. Henry, P. et al. In situpopulation structure andex siturepresentation of the endangered Amur tiger. Molecular Ecology 18, 3173–3184 (2009).

136. Cho, Y. S. et al. The tiger genome and comparative analysis with lion and snow leopard genomes. Nature Communications 4, (2013).

137. Liao, W. & Reed, D. H. Inbreeding-environment interactions increase extinction risk. Animal Conservation 12, 54–61 (2009).

138. Ning, Y. et al. Dispersal of Amur tiger from spatial distribution and genetics within the eastern Changbai mountain of China. Ecology and Evolution 9, 2415–2424 (2019).

139. Howell, L. G. et al. Integrating biobanking minimises inbreeding and produces significant cost benefits for a threatened frog captive breeding programme. Conservation Letters 14, (2020).

140. Bernt, M., Braband, A., Schierwater, B. & Stadler, P. F. Genetic aspects of mitochondrial genome evolution. Molecular Phylogenetics and Evolution 69, 328–338 (2013).

141. Birky, C. W. Uniparental inheritance of organelle genes. Current Biology 18, R692–R695 (2008).

142. Goremykin, V. V., Salamini, F., Velasco, R. & Viola, R. Mitochondrial DNA of Vitis vinifera and the Issue of Rampant Horizontal Gene Transfer. Molecular Biology and Evolution 26, 99–110 (2008).

143. Folk, R. A., Mandel, J. R. & Freudenstein, J. V. Ancestral Gene Flow and Parallel Organellar Genome Capture Result in Extreme Phylogenomic Discord in a Lineage of Angiosperms. Systematic Biology syw083 (2016) doi:10.1093/sysbio/syw083.

144. Yu, H. et al. Genomic evidence for the Chinese mountain cat as a wildcat conspecific (Felis silvestris bieti) and its introgression to domestic cats. Science Advances 7, (2021).

145. Le Roux, J. J., Foxcroft, L. C., Herbst, M. & MacFadyen, S. Genetic analysis shows low levels of hybridization between A frican wildcats (*Felis silvestris lybica*) and domestic cats (*F. s. catus*) in S outh A frica. Ecology and Evolution 5, 288–299 (2014).

146. Pierpaoli, M. et al. Genetic distinction of wildcat (*Felis silvestris*) populations in Europe, and hybridization with domestic cats in Hungary. Molecular Ecology 12, 2585–2598 (2003).

147. Asase, A., Mzumara-Gawa, T. I., Owino, J. O., Peterson, A. T. & Saupe, E. Replacing “parachute science” with “global science” in ecology and conservation biology. Conservation Science and Practice 4, (2021).

148. Stefanoudis, P. V. et al. Turning the tide of parachute science. Current Biology 31, R184–R185 (2021).

149. Li, F.-W. Decolonizing botanical genomics. Nature Plants 7, 1542–1543 (2021).

