## Supplementary figures and images for "Chromosome-level, nanopore-only genome and allele-specific DNA methylation of Pallas’s cat, *Otocolobus manul*"

### Supplemental Figure 1

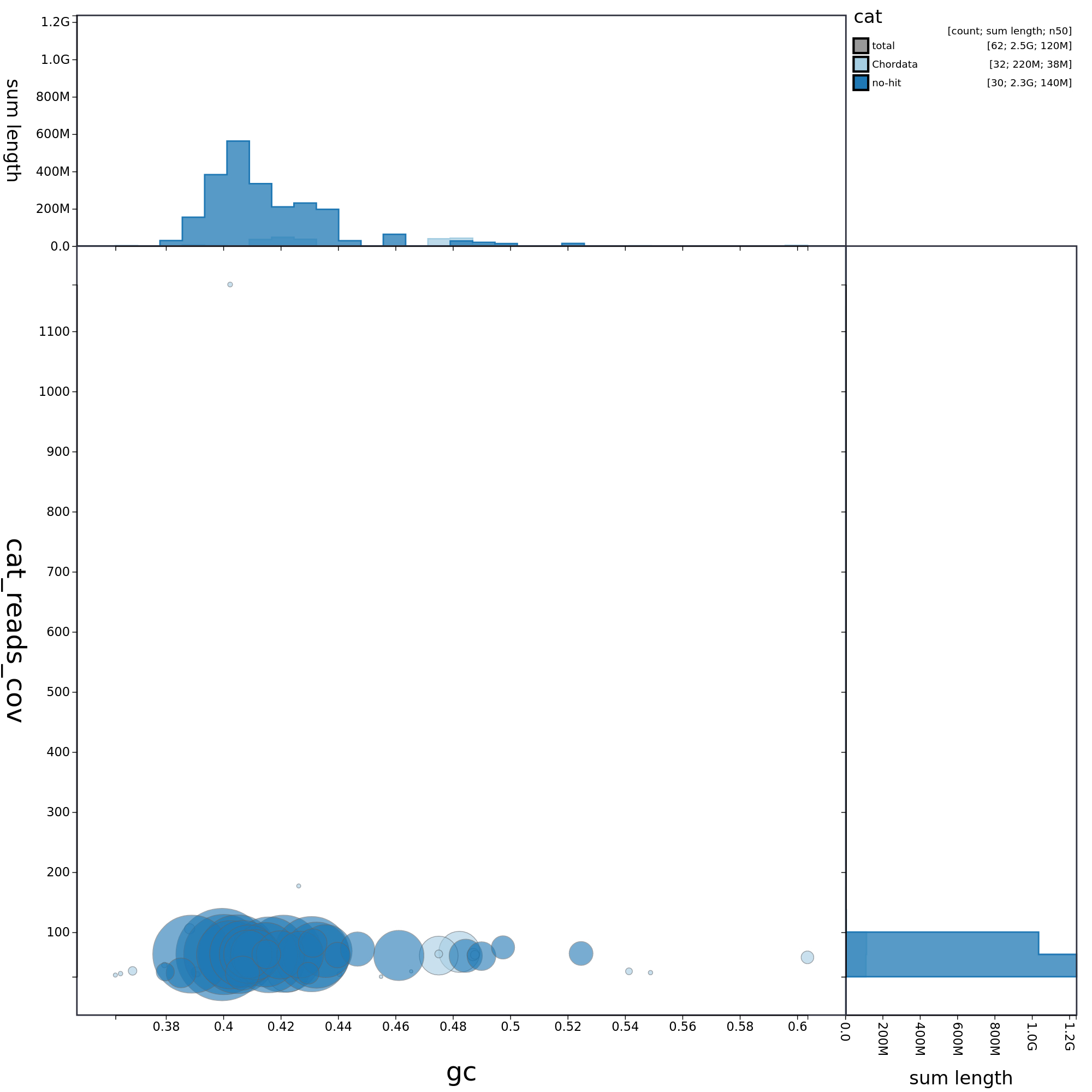

### Supplemental Figure 2

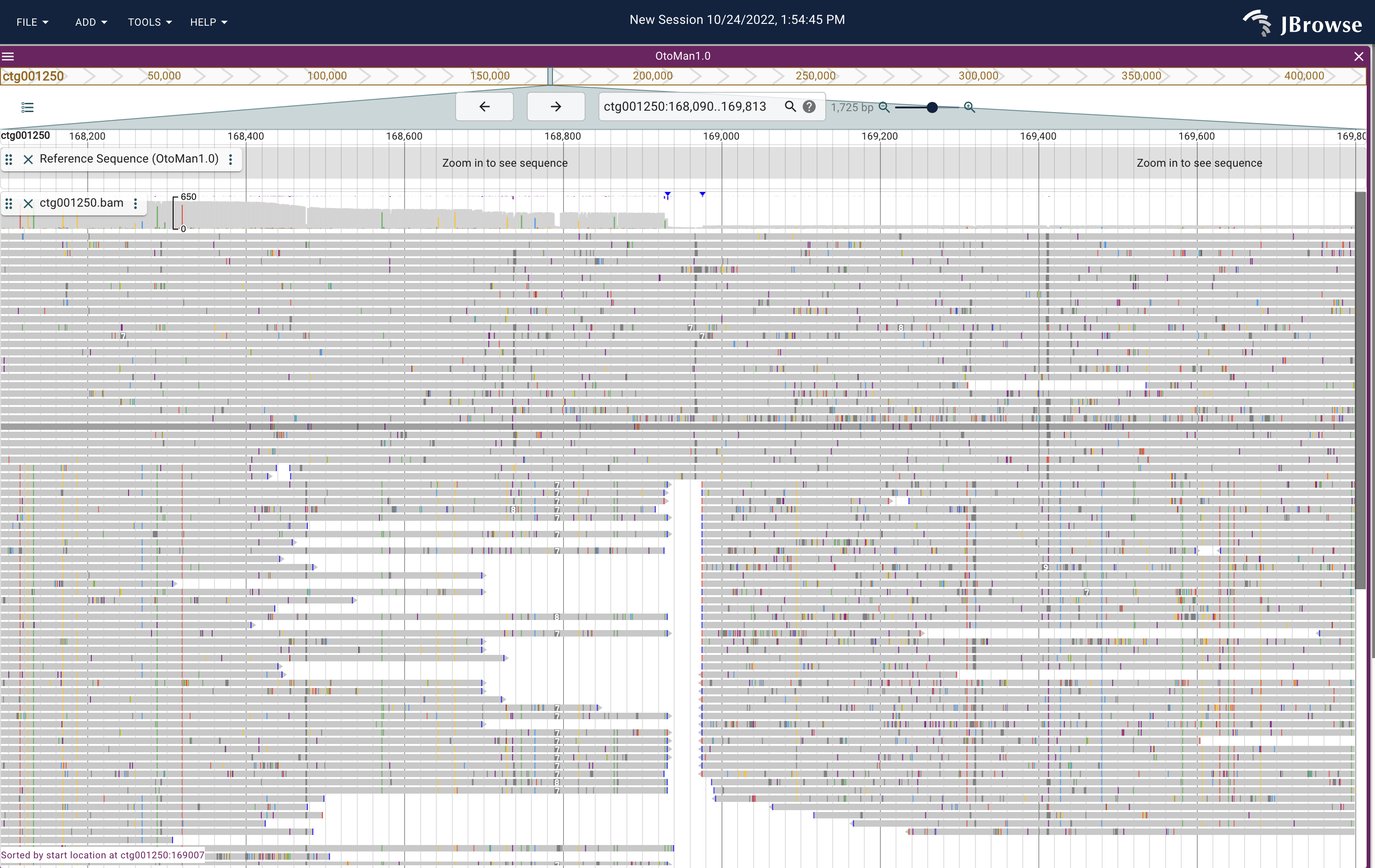

### Supplemental Figure 3

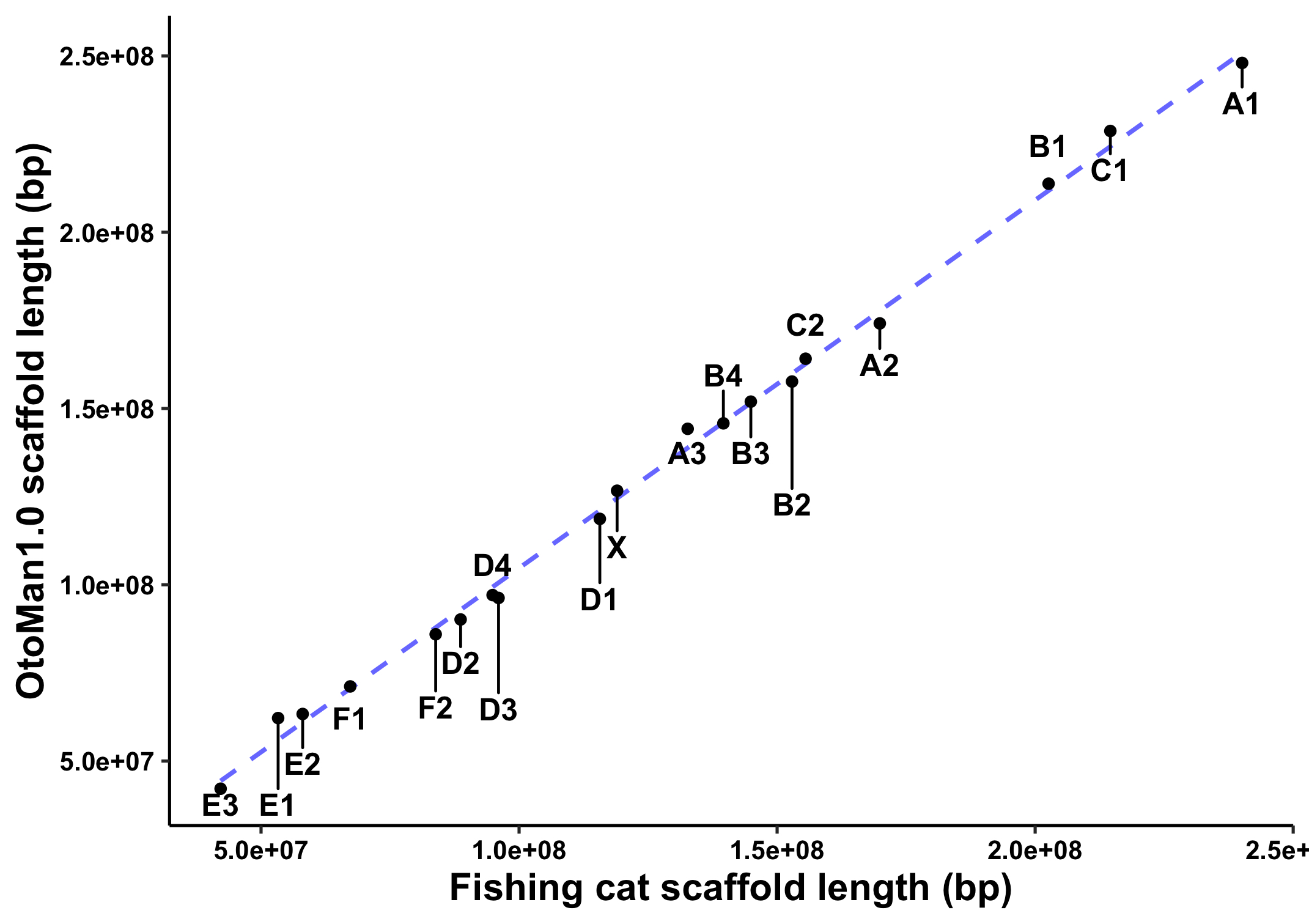

### Supplemental Figure 4

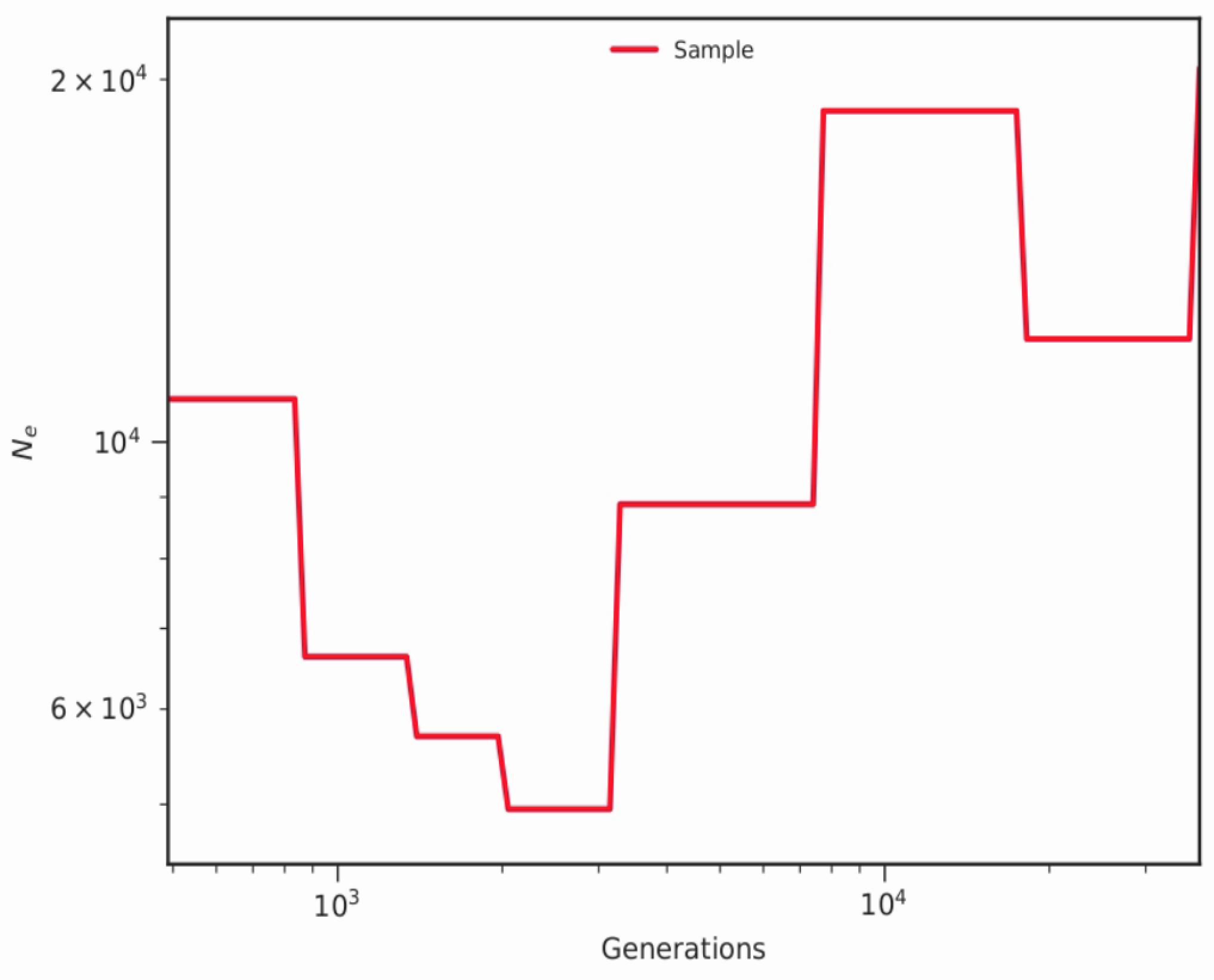
