## Supplemental File 2 for "Chromosome-level, nanopore-only genome and allele-specific DNA methylation of Pallas’s cat, *Otocolobus manul*": Supplemental_File_2.html

Otocolobus manul Nanopore genome assembly


Code 

- Show All Code
- Hide All Code
- Download Rmd

### *Otocolobus manul* Nanopore genome assembly

##### Supplemental Computational Methods

#### Background

A diploid nuclear reference assembly and mitogenome were generated
for Pallas’s cat, *Otocolobus manul,* using a whole blood sample
from a 5-year-old, captive-bred male animal. Allele-specific DNA
methylation analysis, gene annotation, repeat masking, and quality
assessments were also performed.

##### Experimental Design

- Sample origin: Utica Zoo, Utica, NY, USA
- Sample type: Whole blood + DNA Shield (Zymo Research)
- DNA extraction: Zymo Quick DNA Miniprep Plus kit (Zymo
  Research)
- Library prep: ONT SQK-LSK110 kit (Oxford Nanopore)
- Sequencing instrument: MinION flow cell R9.4.1 (Oxford
  Nanopore)
- Sequencing method: Sequencing occurred over 21 days on 7 flow
  cells; 3 libraries were loaded on each flow cell and sequenced for 24
  hours with nuclease flushes in between runs (Wash Kit 004, Oxford
  Nanopore).

##### Computational Resources

The sequencing computer ran Linux Ubuntu 20.04.4 LTS with an AMD
Ryzen 3900x 12-core, 24-thread CPU, 64 GB RAM, and 4 TB SSD storage.
GPU-enabled basecalling was run on two cards, an NVIDIA GeForce 3080Ti
and a 2080Ti. GPU statuses were monitored with `nvtop`.

Downstream assembly analyses and re-basecalling for DNA methylation
were primarily performed on a machine running LInux Ubuntu 20.04.4 LTS
with an AMD Ryzen 9 5950x 16-core, 32-thread CPU, NVIDIA GeForce RTX
3090 GPU, 128 GB RAM, 2 TB SSD storage, and 16 TB NAS storage.
Generation of the final assembly required high-powered cluster computing
resources, primarily due to memory requirements (>100 GB RAM);
statistics for draft assemblies including computational resources and
software calls are available in Table 1 of this document and
Supplemental File 3 of the manuscript.

##### Run length

Seven flow cells generated 15.3 million reads and 158.2 billion bases
in 21 days.

##### Read generation

We used GPU-enabled Guppy v6.2.1 to generate FAST5 files with fast
mode live basecalling. Post-hoc basecalling was performed with the super
accuracy model:

`guppy_basecaller --config dna_r9.4.1_450bps_sup.cfg --compress_fastq --device cuda:0 --nested_output_folder --recursive --input_path no_sample/ --save_path ./sup`

##### Sequencing QC

Nanoq was run with
`nanoq -v -s -i cattotal.fastq.gz`.

#### Assembly and Analysis Methods

##### Draft assembly statistics

Table 1: *O. manul* draft assembly statistics and
computational resources utilized. MSI = Minnesota Supercomputing
Institute, Pegasus = Miami computing cluster.

|  |  |  |  |  |  |  |  |  |  |  |  |
| --- | --- | --- | --- | --- | --- | --- | --- | --- | --- | --- | --- |
| **Name** | **Size** | **Contigs** | **Contig N50** | **Busco** | **Busco carnivora\_odb10** | **Coverage** | **Busco version** | **Assembler** | **CPU time** | **Machine** | **Assembler notes** |
| Shasta min1000 | 2,470,125,084 | 714 | 26,447,788 | 93.9 | C:93.9%[S:92.8%,D:1.1%],F:1.3%,M:4.8%,n:14502 | NA | 5.3.2 | Shasta 0.8.0 | 60 hours | 24 core, 64 Gb RAM | ./shasta-Linux-0.8.0 –input cattotal.fastq –config Nanopore-Oct2021 –memoryBacking disk –memoryMode filesystem –Reads.minReadLength 1000 –assemblyDirectory ShastaRunOct2021 |
| Shasta default | 2,486,571,985 | 846 | 29,762,361 | 94.1 | C:94.1%[S:93.0%,D:1.1%],F:1.4%,M:4.5%,n:14502 | NA | 5.3.2 | Shasta 0.8.0 | 17 hours | 32 core, 128 Gb RAM | ./shasta-Linux-0.8.0 –input cattotal.fastq –config Nanopore-Oct2021 –memoryBacking disk –memoryMode filesystem –assemblyDirectory ShastaRunOct2021 |
| Shasta default +racon1x | 2,444,972,901 | 512 | 26,196,195 | 83.1 | C:83.1%[S:82.3%,D:0.8%],F:2.0%,M:14.9%,n:14502 | NA | 5.3.2 | Shasta 0.8.0 | 21 hours | 24 core, 64 Gb RAM | ./shasta-Linux-0.8.0 –input cattotal.fastq –config Nanopore-Oct2021 –memoryBacking disk –memoryMode filesystem –assemblyDirectory ShastaRunOct2021 |
| Shasta default +medaka | 2,472,619,612 | 714 | 26,472,672 | 94.1 | C:94.1%[S:93.0%,D:1.1%],F:1.2%,M:4.7%,n:14502 | NA | 5.3.2 | Shasta 0.8.0 |  |  | medaka\_consensus -b 100 -i ../cattotal.fastq -d Assembly.fasta -o medaka-results/ -m r941\_min\_sup\_g507 -t 23 |
| Flye over10k | 2,497,029,119 | 850 | 21,167,348 | 93.5 | C:93.5%[S:92.3%,D:1.2%],F:1.8%,M:4.7%,n:14502 | 41x | 5.3.2 | Flye 2.9-b1774 | 14 hours | MSI 128 core, 2 TB RAM | Flye/bin/flye –nano-hq catOver10k.fastq.gz –out-dir flye\_over30k –threads 128 |
| Flye over30k\_asm | 2,491,940,851 | 816 | 16,256,589 | 91.5 | C:91.5%[S:90.4%,D:1.1%],F:2.3%,M:6.2%,n:14502 | 11x | 5.3.2 | Flye 2.9-b1774 | 10.5 hours | 32 core, 128 Gb RAM | flye –nano-hq catOver30k.fastq.gz –out-dir flye\_output –genome-size 3g –threads 28 –asm-coverage 40 |
| Flye over30k\_asm\_polish-target | 2,491,762,986 | 815 | 16,256,965 | 93.4 | C:93.4%[S:92.2%,D:1.2%],F:1.8%,M:4.8%,n:14502 | 64x | 5.3.2 | Flye 2.9-b1774 | 13 hours | 32 core 128 G RAM | flye –nano-hq ../cattotal.fastq –polish-target assembly.fasta -o cattotal\_polish\_target-1 –threads 14 |
| Flye over30k | 2,491,811,451 | 851 | 16,585,934 | 91.4 | C:91.4%[S:90.4%,D:1.0%],F:2.3%,M:6.3%,n:14502 | 11x | 5.3.2 | Flye 2.9-b1774 | 4 hours | MSI 128 core, 2 TB RAM | Flye/bin/flye –nano-hq catOver30k.fastq.gz –out-dir flye\_results\_over30k –genome-size 3g –threads 128 |
| Flye best15g | 2,297,972,615 | 4316 | 1,321,834 | 66.5 | C:66.5%[S:65.7%,D:0.8%],F:2.7%,M:30.8%,n:14502 | 6x | 5.3.2 | Flye 2.9-b1768 | 14 hours | Pegasus 8 nodes, 1500MB RAM per core | flye –nano-hq catbest15g.fastq –out-dir cat15\_flye\_assembly –threads 15 |
| Flye best15g HQ medaka | 2,299,902,380 | 4316 | 1,323,637 | 74.2 | C:74.2%[S:73.2%,D:1.0%],F:2.0%,M:23.8%,n:14502 |  | 5.3.2 | Medaka 1.6.0 Flye 2.9-b1768 | 48 hours | Pegasus 6 nodes, 1500MB RAM per core | medaka\_consensus -b 100 -i cattotal.fastq -d cat15\_flye\_assembly.fasta -t 6 -m r941\_min\_sup\_g507 |
| Flye best79g | 2,492,484,939 | 650 | 16,872,056 | 93.6 | C:93.6%[S:92.5%,D:1.1%],F:1.7%,M:4.7%,n:14502 | 30x | 5.3.2 | Flye 2.9-b1768 | 30 hours | Pegasus 15 nodes, 1500MB RAM per core | flye –nano-hq catbest79g.fastq –out-dir cat79\_flye\_assembly –threads 10 |
| Flye best79g HQ medaka | 2,495,464,699 | 650 | 16,894,862 | 94.7 | C:94.7%[S:93.5%,D:1.2%],F:1.2%,M:4.1%,n:14502 |  | 5.3.2 | Medaka 1.6.0 Flye 2.9-b1768 | 48 hours | Pegasus 6 nodes, 1500MB RAM per core | medaka\_consensus -b 100 -i cattotal.fastq -d cat79\_flye\_assembly.fasta -t 6 -m r941\_min\_sup\_g507 |
| Flye cattotal\_asm | 2,489,000,215 | 1046 | 28,590,467 | 93.6 | C:93.6%[S:92.4%,D:1.2%],F:1.7%,M:4.7%,n:14502 | 61x | 5.3.2 | Flye 2.9-b1768 | 15 hours | MSI 128 core, 2 TB RAM | Flye/bin/flye –nano-hq cattotal.fastq.gz –out-dir flye\_results\_cattotal –genome-size 2.5g –asm-coverage 40 –threads 128 |
| Flye cattotal\_asm+3xpolish-target+medaka | 2,491,830,032 | 1041 | 28,623,091 | 94.8 | 94.8%[S:93.6%,D:1.2%],F:1.2%,M:4.0%,n:14502 |  | 5.3.2 | Flye 2.9-b1768 Medaka 1.6.0 | 72 hours | 16 core, 128 G RAM |  |
| Flye cattotal | 2,490,902,702 | 1139 | 27,172,101 | 93.5 | C:93.5%[S:92.3%,D:1.2%],F:1.7%,M:4.8%,n:14502 | NA | 5.3.2 | Flye 2.9-b1768 | 15.5 hours | MSI 128 core, 2 TB RAM | Flye/bin/flye –nano-hq cattotal.fastq.gz –out-dir flye\_results\_cattotal –genome-size 2.5g –threads 128 |
| Flye cattotal\_asm+medaka+nextpolish | 2,495,697,703 | 1046 | 28,662,605 | 94.5 | C:94.5%[S:93.3%,D:1.2%],F:1.3%,M:4.2%,n:14502 |  | 5.3.2 | NextPolish 1.4.0 | 2 hours | 32 core, 128 G RAM |  |
| Flye cattotal\_asm+medaka | 2,492,512,061 | 1046 | 28,628,316 | 94.8 | C:94.8%[S:93.6%,D:1.2%],F:1.2%,M:4.0%,n:14502 | NA | 5.3.2 | Medaka 1.6.0 Flye 2.9-b1768 | 11 hours | 32 core, 128 Gb RAM | medaka\_consensus -b 100 -i cattotal.fastq -d cattotal\_assembly\_flye.fasta -o medaka\_flye\_cattotal/ -t 28 -m r941\_min\_sup\_g507 |

##### Generation of final assembly

The two highest-scoring assemblies were from Flye and NextDenovo,
respectively, each using the full readset.

The Flye assembly was generated on the UMN computing cluster due to
memory requirements (2 TB RAM). Polishing was performed locally with Medaka.

```
# Assemble
flye --nano-hq cattotal.fastq.gz --out-dir flye_results_cattotal --genome-size 2.5g --asm-coverage 40 --threads 128

# Polish
medaka_consensus -b 100 -i cattotal.fastq -d cattotal_assembly_flye.fasta -o medaka_flye_cattotal/ -t 28 -m r941_min_sup_g507
```

The NextDenovo assembly was generated and polished locally.

```
# Contents of run.cfg file
[General]
job_type = local # local, slurm, sge, pbs, lsf
job_prefix = nextDenovo
task = all # all, correct, assemble
rewrite = yes # yes/no
deltmp = yes 
parallel_jobs = 1 # number of tasks used to run in parallel
input_type = corrected # raw, corrected
read_type = ont # clr, ont, hifi
input_fofn = input.fofn
workdir = 01_rundir

[correct_option]
read_cutoff = 1k
genome_size = 2.4g # estimated genome size
sort_options = -m 20g -t 15
minimap2_options_raw = -t 23
pa_correction = 1 # number of corrected tasks used to run in parallel, each corrected task requires ~TOTAL_INPUT_BASES/4 bytes of memory usage.
correction_options = -p 15

[assemble_option]
minimap2_options_cns = -t 23 
nextgraph_options = -a 1

# Assemble
nohup ./NextDenovo/nextDenovo run.cfg &

# Polish
medaka_consensus -b 100 -i cattotal.fastq -d nd.asm.fasta -o medaka-results/ -m r941_min_sup_g507
```

The Flye and NextDenovo assemblies were combined using the Quickmerge
metassembler to increase contiguity while preserving quality.

```
# Align assemblies to be merged with Nucmer twice, alternating which is reference and which is query
nucmer --threads 23 --prefix flye-nd nd_medaka_consensus.fasta medaka_flye_cattotal_consensus.fasta

nucmer --threads 23 --prefix nd-flye medaka_flye_cattotal_consensus.fasta nd_medaka_consensus.fasta

# Run Quickmerge twice, keeping whichever version has best contiguity + BUSCO, then re-merge that version with the original NextDenovo + Medaka assembly. One run is shown.
quickmerge -d out.rq.delta -q nd_medaka_consensus.fasta -r medaka_flye_cattotal_consensus.fasta -hco 5.0 -c 1.5 -l 26472672 -ml 10000 -p prefix
```

Since BUSCO duplicates were reported as >1%, we ran the Purge Haplotigs pipeline
for reassignment and removal of allelic contigs.

```
# align reads to output of Quickmerge
minimap2 -t 20 -ax map-ont ../final_assembly/merged_3waymerge.fasta ../cattotal.fastq | samtools sort -m 1G -o aligned.bam -T tmp.ali

# Coverage histogram
purge_haplotigs  hist  -b aligned.bam  -g ../medaka-results/consensus.fasta  -t 20

# There were two peaks at 30x and 60x with a trough at 40x. 

# Analyze per-contig coverage
purge_haplotigs cov -i aligned.bam.gencov -l 15 -m 40 -h 100 -o coverage_stats.csv

# Run the purging pipeline
purge_haplotigs purge -g ../final_assembly/merged_3waymerge.fasta -c coverage_stats.csv -t 20 -d -b aligned.bam
```

The resulting `curated.fasta` went from 99 to 62 sequences
with only very minimal reduction in length and N50.

##### Sequencing contamination

After realignment of sequencing reads with Minimap2, raw read
contamination was assessed using Kraken2, BLAST, and Pavian.

Kraken2 was used with the NCBI nt database for microbial DNA
detection in the unmapped reads.

```
# Filter for unmapped reads
samtools view -f 4 curated.bam > unmapped.curated.sam

# Convert from SAM to BAM to FASTQ
samtools view -b unmapped.curated.sam > unmapped.curated.bam

samtools fastq unmapped.curated.bam > unmapped.consensus.fastq

# Run Kraken2
kraken2 --db /mnt/bc91d872-d479-41b6-b994-c521c27d41cf/kraken2/k2_pluspf16gb/ --threads 10 --use-names --report cattotal.FASTQ.unmapped.report.txt --output cattotal.FASTQ.unmapped.out.txt unmapped.consensus.fastq

# Visualize with Pavian - From inside R type:
pavian::runApp(port=5000)
```

To run local BLAST, first downloaded the nt database from NCBI
via FTP.

Then ran megablast:

```
blastn -query consensus.fasta -task megablast -db ~/Desktop/genomes/nt/nt -outfmt '6 qseqid staxids bitscore std sscinames sskingdoms stitle' -culling_limit 10 -num_threads 23 -evalue 1e-3 -out consensus.fasta.vs.nt.cul5.1e3.megablast.out
```

Anomalous contig GC content and coverage (<1x or >1000x) were
assessed with Blobtools2.
Downloaded taxdump from NCBI,
unzipped it, and moved all the resulting `.dmp` files to the
subdirectory `taxdump`.

```
# Re-align reads to assembly with minimap2
minimap2 -ax map-ont -t 28 curated.wrapped.fasta cattotal.fastq.gz | samtools view -F 4 -h -@ 28 | samtools sort -@ 28 -O BAM > curated_cattotal.sorted.bam

samtools index curated.wrapped.sorted.bam

# Create blobdir
 blobtools create \
 --fasta data/curated.wrapped.fasta \
 --meta data/cat.yaml \
 --taxid 61408 \
 --taxdump taxdump/ \
 datasets/cat
 
# Add BLAST results to blobdir
blobtools add --hits data/partial.combined.curated.vs.nt.cul5.1e3.megablast.out --taxdump taxdump/ datasets/cat

# Generate coverage file with SAMtools
samtools coverage curated_cattotal.sorted.bam > curated.coverage.txt

# Add coverage file to blobdir
blobtools add --text data/curated.coverage.txt --text-header --text-cols '#rname=identifier,meandepth=cat_reads_cov' datasets/cat

# Fix plot axes
blobtools add --key plot.y=cat_reads_cov datasets/cat

# Run BUSCO for assembly completeness
busco -i curated.wrapped.fasta -o curated_busco -m genome -l carnivora_odb10 -c 28

# Add BUSCO results to blobdir
blobtools add --busco data/full_table.tsv datasets/cat

# View blobdir interactively
blobtools view --local --interactive datasets/cat
```

A 420 kb contig with >1000x coverage and no
*Carnivora*-specific BUSCO genes matching
*Felidae* by identity, ctg001250, was removed to generate the
final primary assembly.

`seqkit grep -v -p ctg001250 curated.wrapped.fasta > no_ctg001250_curated.wrapped.fasta`

BUSCO was re-run and reads were re-aligned with Minimap2 as described
above.

##### Coverage and quality statistics

Assembly coverage was assessed using Mosdepth.

`mosdepth -n --fast-mode -t 8 no_ctg001250_curated no_ctg001250_curated.cattotal.sorted.bam`

Reference-free, k-mer-based quality statistics were generated with Merqury.

```
# In theory, merfin can create the QV score but in practice it requires more memory than available (>65 Gb for Pallas's cat). Merqury uses the same meryl k-mer database but does not require so much memory and completes within minutes.

# Install merfin which will install a good version of meryl
git clone https://github.com/arangrhie/merfin.git
cd merfin/src
make -j 12

# Construct k-mer db
meryl count k=$k reads.fastq.gz output reads.meryl

# Collect histogram for GenomeScope 2.0 which makes nice graphics
meryl histogram reads.meryl > reads.hist

# Exclude frequency = 1 k-mers
meryl greater-than 1 reads.meryl output reads.gt1.meryl

# Install merqury via conda
conda create -n merqury -c conda-forge -c bioconda merqury
conda activate merqury

# Two assemblies (diploid), so no hap-mers
$MERQURY/merqury.sh read-db.meryl asm1.fasta asm2.fasta out_prefix
```

##### Repeat identification

Genomic repeat content was assessed with RepeatMasker.

```
RepeatMasker curated.fasta -s -pa 16 -species Carnivora -xsmall
```

The multi-cat repeat plot was built by locally running RepeatMasker
on felid genomes and loading the results from a manually organized
spreadsheet, which is included in Supplemental File 3 of the
manuscript.

```
library(tidyverse) library(ggpubr)

cat_repeats_df <- readRDS("CatRepeatsDF.Rds")

cat_repeats_tibble <- readRDS("RepeatTibble.Rds") %>% 
                      mutate(Element = 
                        str_replace(Element, "unclassified", "Unclassified"),
                        Element = str_replace(Element, "DNA elements", "DNA Elements"), 
                        Element = str_replace(Element, "LTR", "LTRs"))

cat_repeats_tibble$Element <- factor(cat_repeats_tibble$Element, 
                              levels = rev(c( "LINEs", "SINEs", "LTRs",
                              "Simple Repeats", "Low Complexity", 
                              "DNA Elements", "Small RNA", "Unclassified" )))

colorblind_hex <- c('#0173b2', '#de8f05', '#029e73', '#d55e00', '#cc78bc', '#ca9161', '#fbafe4', '#949494')

cat_repeat_plot <- cat_repeats_tibble %>% 
                        ggplot(aes(x = Species, y = PercentSequence, 
                          fill = Element)) + 
                        geom_col(position = position_stack()) + 
                        labs(y = "Percent of genome sequence") + 
                        theme_pubr() + labs_pubr() + 
                        theme(axis.text.x = 
                          element_text(angle = 45, hjust = 1)) +
                        scale_fill_manual(values = rev(colorblind_hex)) +
                        theme(legend.title = 
                          element_blank(), legend.justification = "center") +
                        guides(fill=guide_legend(nrow=2,byrow=T, reverse = T))

ggsave("cat_repeats.png", plot = cat_repeat_plot, dpi = "retina", width = 18, height = 15, unit = "cm")
```

##### Gene annotation

GeMoMa v1.8 was
applied to the primary assembly for homology-based protein prediction
with the *F. catus* gene annotation (Fca126\_mat1.0;
GCF\_018350175.1) used as the reference.

```
java -Xmx50g -jar GeMoMa-1.8.jar CLI GeMoMaPipeline \
  threads=28 outdir=Fcat_annotation \
  GeMoMa.Score=ReAlign AnnotationFinalizer.r=NO \
  o=true t=no_ctg001250_curated.wrapped.fasta \
  i=F_catus a=GCF_018350175.1_F.catus_Fca126_mat1.0_genomic.gff.gz \
  g=GCF_018350175.1_F.catus_Fca126_mat1.0_genomic.fna.gz
```

Protein models were scored using BUSCO’s protein mode.

`busco -i predicted_proteins.fasta -o busco -m protein -l carnivora_odb10 -c 28`

##### Variant Calling

Variant calling and phasing was performed using the PEPPER-Margin-DeepVariant
pipeline.

```
sudo docker pull kishwars/pepper_deepvariant:r0.8

sudo docker run \
-v "/home/niki/Documents/pallas_cat/chris_curated_assembly/data":"/home/niki/Documents/pallas_cat/chris_curated_assembly/data" \
-v "/home/niki/Documents/pallas_cat/chris_curated_assembly/data/pepper_output":"/home/niki/Documents/pallas_cat/chris_curated_assembly/data/pepper_output" \
kishwars/pepper_deepvariant:r0.8 \
run_pepper_margin_deepvariant call_variant \
-b /home/niki/Documents/pallas_cat/chris_curated_assembly/data/no_ctg001250_curated.cattotal.sorted.bam \
-f /home/niki/Documents/pallas_cat/chris_curated_assembly/data/no_ctg001250_curated.wrapped.fasta \
-o /home/niki/Documents/pallas_cat/chris_curated_assembly/data/pepper_output \
-t 28 \
--ont_r9_guppy5_sup \
--phased_output
```

Variant statistics were generated using Whatshap version 1.4 and
VCFtools version
0.1.17.

`whatshap stats PEPPER_MARGIN_DEEPVARIANT_FINAL_OUTPUT.phased.vcf.gz --tsv=pepper_phased_whatshap.tsv`

`vcftools --TsTv-summary --gzvcf mod.phased.vcf.gz`

A consensus FASTA for the secondary haplotype was generated by
applying all biallelic variants to the final assembly (considered the
primary haplotype) with the consensus module of BCFtools version
1.15.1.

```
cat no_ctg001250_curated.wrapped.fasta | bcftools consensus \
PEPPER_MARGIN_DEEPVARIEANT_FINAL_OUTPUT.phased.vcf.gz > haplotype2_noctg001250_consensus.fasta
```

Runs of homozygosity were assessed with SMC++.

```
sudo docker run --rm -v $PWD:/mnt terhorst/smcpp:latest vcf2smc PEPPER_MARGIN_DEEPVARIANT_FINAL_OUTPUT.phased.vcf.gz out/ctg001740.smc.gz ctg001740 Sample:Sample

# Here, 0.86e-8 represents the mammalian mutation rate per year.
sudo docker run --rm -v $PWD:/mnt terhorst/smcpp:latest estimate -o analysis/ 0.86e-8 out/ctg001740.smc.gz

sudo docker run --rm -v $PWD:/mnt terhorst/smcpp:latest plot plot.pdf analysis/model.final.json 

# The first plot command gives plots in generations
sudo docker run --rm -v $PWD:/mnt terhorst/smcpp:latest plot plot.pdf analysis/model.final.json 

# The second command gives plot in years assuming generation time is 3 years.
sudo docker run --rm -v $PWD:/mnt terhorst/smcpp:latest plot plot.pdf analysis/model.final.json -g 3
```

##### DNA Methylation

DNA methylation (5mC) at cytosine guanine dinucleotides (CpGs) was
determined by re-basecalling QC-passed FAST5 files with a modified base
configuration of GPU-mode Guppy (dna\_r9.4.1\_450bps\_modbases\_5mc\_cg\_sup).
The final primary assembly was used as a reference.

```
~/Desktop/ont-guppy/bin/guppy_basecaller \
--input_path fast5_pass \
-s aln_modbams \
-c dna_r9.4.1_450bps_modbases_5mc_cg_sup.cfg \
--bam_out --align_ref no_ctg001250_curated.wrapped.fasta \
-x "cuda:0"
```

The resulting modified BAMs (modBAMs) for each flow cell directory
were concatenated together into a single file, then sorted and indexed
with SAMtools.

Cat together modbams for each flow cell directory:

`samtools cat -o cat1_aln_mod.bam pass/*.bam -@ 20`

Then `mv`ed all those files up a few directories to
`cat-analysis` to cat them all together into one total
aligned modbam, `aln_cattotal_mod.bam`.

Sorted with
`samtools sort -o aln_cattotal_mod.sorted.bam -@ 28 aln_cattotal_mod.bam`

Indexed with
`samtools index -@ 28 aln_cattotal_mod.sorted.bam`

PEPPER-Margin-DeepVariant was then re-run to generate a haplotagged
modBAM.

```
sudo docker run \
-v "/media/niki/athena_storage/pallas_cat":"/media/niki/athena_storage/pallas_cat" \
-v "/mnt/synology/cat-analysis/aln_modbams":"/mnt/synology/cat-analysis/aln_modbams" \
kishwars/pepper_deepvariant:r0.8 \
run_pepper_margin_deepvariant call_variant \
-b /mnt/synology/cat-analysis/aln_modbams/aln_cattotal_mod.sorted.bam \
-f /media/niki/athena_storage/pallas_cat/no_ctg001250_curated.wrapped.fasta \
-o /media/niki/athena_storage/pallas_cat \
-p mod \
-t 28 \
--ont_r9_guppy5_sup \
--phased_output
```

This file and the primary assembly were used as input for modbam2bed, which
aggregates modified base counts to generate bedMethyl files.

```
modbam2bed -m 5mC -t 28 -e -d 1000 --aggregate --cpg \
--haplotype=1 --prefix=no_ctg001250_HP1 \
no_ctg001250_curated.wrapped.fasta \
pepper_modbam/mod.haplotagged.bam > modbam_2bed/no_ctg001250_mod.HP1.bed
```

Switched `--haplotype=1` and `HP1` to
`HP2` to get bed file for haplotype 2. Modified the files to
meet DSS input requirements with:

```
awk -v OFS='\t' '{print $1, $2, $12+$13, $13}' \
no_ctg001250_mod.HP1.bed > no_ctg001250_mod.HP1.dss
```

Ran modbam2bed without haplotype flag to get bedMethyl file for
calculating global methylation on the collapsed data.

```
modbam2bed -m 5mC -t 28 -e -d 1000 --aggregate --cpg \
--prefix=no_ctg001250_collapsed \
no_ctg001250_curated.wrapped.fasta \
pepper_modbam/mod.haplotagged.bam > modbam_2bed/no_ctg001250_mod.collapsed.bed
```

Then ran
`awk '$5>0 && $12+$13>10 {total+=$11; count++} END{print total/count}'`
on the output to get average percent methylation.

Allele-specific DNA methylation was analyzed using the R package DSS
developer version 2.43.2, which has improved parallelization capacity.
The package’s two-group statistical comparison module
`DMLtest` was used to identify differentially methylated loci
between the two haplotypes. The `callDMR` module was then
used to identify multi-CpG differentially methylated regions (DMRs).

```
library(DSS)
library(tidyverse)

setwd("/media/niki/athena_storage/pallas_cat/modbam_2bed")

dss_cols <- c("chr", "pos", "N", "X")

hp_1 <- read_tsv("no_ctg001250_mod.HP1.dss",
                 show_col_types = F,
                 col_names = dss_cols)

hp_2 <- read_tsv("no_ctg001250_mod.HP2.dss",
                 show_col_types = F,
                 col_names = dss_cols)

hap_list <- list(HP1 = hp_1, HP2 = hp_2)

rm(hp_1, hp_2)

cov_filter <- function(x) {
  y <-  x %>%
    filter(N >= 10)
  return(y)
}

hap_list_filtered <- map(hap_list, cov_filter)

bsobj <- makeBSseqData(hap_list_filtered,
                       names(hap_list_filtered))

dmlTest <- DMLtest(
  bsobj,
  group1 = "HP2",
  group2 = "HP1",
  smoothing = T,
  smoothing.span = 500,
  ncores = 28
)

dmrs <- callDMR(dmlTest, 
                p.threshold = 0.001,
                delta = 0.5,
                minlen = 100,
                minCG = 15,
                dis.merge = 1500
                )
```

To circumvent the relatively low BUSCO quality of our gene
annotation, DMR sequences were annotated via lifting of Felis catus
reference (GCF\_018350175.1) gene features onto the O. manul assembly
using Liftoff. The nearest feature to each DMR was then identified using
AGAT and the
`closest` module of BEDTools. DMRs
annotated near genes with a “LOC” symbol were additionally annotated
with an alias or gene description, when available, via manual look up in
NCBI’s Gene database.

```
# GTF conversion with AGAT
agat_convert_sp_gxf2gxf.pl --gff Fcat_contig_liftoff.sorted.gtf -o Fcat_contig_liftoff.sorted.gff

# Filter for gene features
cat Fcat_contig_liftoff.sorted.gff | grep -v '#' | \
  awk 'BEGIN{FS=OFS="\t"}{ if($3=="gene"){print} }' \
  > Fcat_liftoff_gene.gff

# Sort  
bedtools sort -i Fcat_liftoff_gene.gff > Fcat_liftoff_gene.sorted.gff

# Find nearest feature
bedtools closest -d -s -a dmrs.sorted.bed -b Fcat_liftoff_gene.sorted.gff > closest_liftoff_dmr.bed
```

Visualizations were generated using Methylartist
version 1.2.3, which required haplotagging the PEPPER output with Longphase, to validate
each DMR. At the time of our run, Methylartist required phase tags in
the VCF file even if the option to ignore them was included in the
call.

```
# haplotagging PEPPER output with longphase for methylartist input
longphase_linux-x64 haplotag \
-s pepper_modbam/mod.phased.vcf.gz \
-b aln_cattotal_mod.sorted.bam \
-t 28 \
-o longphase_modbam

# Run methylartist
methylartist locus --ref no_ctg001250_curated.wrapped.fasta \
--motif CG --phased --color_by_hp --ignore_ps \
--phase_labels 1:HP1,2:HP2 --samplepalette husl --labelgenes \
-b longphase_haplotagged_pepper_mod.bam --nticks 6 \
--gtf Fcat_contig_liftoff.sorted.gtf.gz \
--statname methyl --highlight_panels 0.2 \
-l ctg000530:65947332-65947927 -i ctg000530:65945332-65949927
```

##### Mitochondrial genome

Nanopore reads were aligned to the *Felis catus* mitogenome
using Minimap2.

```
minimap2 -ax map-ont Fca126.mtDNA.fasta cattotal.fastq -a -o cattotal.vs.Fca126.mtDNA.sam -t 24
```

Sorted and filtered for mapped reads only and convert to BAM:

`samtools sort cattotal.vs.Fca126.mtDNA.sam -o cattotal.vs.Fca126.mtDNA.bam`

`samtools view -F 4 -b cattotal.vs.Fca126.mtDNA.bam > cattotal.vs.Fca126.mtDNA.aln.bam`

`samtools index cattotal.vs.Fca126.mtDNA.aln.bam`

`samtools fastq cattotal.vs.Fca126.mtDNA.aln.bam > cat-mtDNAreads_mapped.fastq`

Aligned reads were then downsampled to 10,000 reads and Flye in
metagenome mode was used for assembly.

`seqtk sample cat-mtDNAreads_mapped.fastq 10000 > cat-mtDNAreads_mapped_subsample.fastq`

```
flye --nano-hq cat-mtDNAreads_mapped_subsample2.fastq \
--out-dir cat_mito_flye_10Kdownsamples_HQassembly \
--genome-size 16k --meta --threads 10
```

Mitogenome annotation and manual rearrangement to start the circular
mitogenome at COX1 was done as in Wanner et al.
(2021):

1. Pulled out circular, mitogenome-sized contig from Flye
   output
2. Uploaded FASTQ file to MITOS2 GUI and
   exported TBL with annotations
3. Used MITOS2 TBL annotations to determine start site of COX1, then
   manually cut from COX1 start and pasted everything before the COX1 start
   to the end:

   - `cat *.fasta | cut -b 6200-` and
     `cat *.fasta | cut -b -6200`
4. Used GALAXY to convert TBL + FASTA to .gb file and visualized in
   Open
   Vector Editor

A mitogenome phylogeny was built using IQ-TREE version 1.6.12 after
multi-alignment with the MUSCLE GUI. Following
IQ-TREE formatting requirements, we prepared the resulting file and
uploaded to the IQ-TREE
webserver.

IQ-TREE was run with the default parameters described here to run
1000 bootstrap maximum likelihood trees:

1. Uploaded sequences
2. Under “bootstrapping” selected “ultrafast” and 1000
   iterations
3. Kept default tree search parameters

The final tree visualization was built with IcyTree, which requires the IQ-TREE
output to be in newick format.

##### Scaffolding

The contig-level assembly was scaffolded onto the fishing cat (*P.
viverrinus*) reference genome with the `scaffold` module
of RagTag. Options were
set to infer gap lengths and a file was provided to exclude chrUns.

```
ragtag.py scaffold \
-e exclude_chrUns_Pviv.txt \
-o ragtag_scaffold_infergaps \
-w -r -t 28 \
--aligner ~/Desktop/mm2-fast/minimap2 \
GCF_022837055.1_UM_Priviv_1.0_genomic.fna \
no_ctg001250_curated.wrapped.fasta
```

The result was assessed with
`seqkit stat -a ragtag.scaffold.fasta` and by examining the
`.stats` output file of RagTag.

Alignment of the two genomes was assessed with Dot, which calls Nucmer
from the package MUMmer4.

```
# align genomes with Nucmer
nucmer -p GCF_022837055.1_UM_Priviv_1.0_genomic.fna no_ctg001250_curated.wrapped.fasta -t 28

# Prep files for visualization
python3 dot/DotPrep.py --delta Pviv_no_ctg001250_curated.delta

# Visualize with Dot browser
firefox dot/index.html # upload associated output.coords and output.coords.idx
```

An ideogram of contig positions in the scaffolds was generated with
the R package chromoMap.

```
# annot file
seqkit locate --bed --pattern-file no_ctg001250_curated.wrapped.fasta \
ragtag_scaffold_infergaps.fasta --threads 28 > ctg_in_scaf.bed

# add 1 to stard because they're zero based and chromomap wants 1 based
awk -F "\t" 'BEGIN{OFS="\t"} {print $4,$1,$2+1,$3}' ctg_in_scaf.bed > ctg_in_scaf_chromomap.bed

# chrom file
seqkit fx2tab -l -g -n -i -H ragtag_scaffold_infergaps.fasta | awk -F "\t" 'BEGIN{OFS="\t"} NR > 1 {print $1,1,$2}' > scaffold_chrs_chromomap.bed
```

Mutated the annotation file to distinguish contigs:

```
library(tidyverse)

df = read_tsv("ctg_in_scaf_chromomap.bed",col_names=c("ctg","chr","start","end"))

df_nest = df %>% group_by(chr) %>% nest() %>% mutate(data2 = map(data, ~ rownames_to_column(.x, var = "ID")))

df_id = df_nest %>% unnest(data2) %>% dplyr::select(-data) %>% filter(chr != "ctg000650_RagTag") %>% # no BUSCOs filter(chr != "ctg000740_RagTag") %>% # no BUSCOs relocate(ID, .after = end) %>% relocate(chr, .after = ctg)

write_tsv(df_id, "id_ctg_scaff_chromomap.bed", col_names = F)
```

Replaced the scaffold names with chromosome names in each file:

`head -n -2 scaffold_chrs_chromomap.bed > full_scaffold_chrs_chromomap.bed`

Edited chromosome names:

```
chr_df <- read_tsv("full_scaffold_chrs_chromomap.bed",
                   col_names = c("chr", "start", "end"))

scaff_names <- tibble::tribble(
  ~ scaffold, ~ chr_name,
  'NC_062561.1_RagTag', 'A1',
  'NC_062562.1_RagTag', 'A2',
  'NC_062563.1_RagTag', 'A3',
  'NC_062564.1_RagTag', 'B1',
  'NC_062565.1_RagTag', 'B2',
  'NC_062566.1_RagTag', 'B3',
  'NC_062567.1_RagTag', 'B4',
  'NC_062568.1_RagTag', 'C1',
  'NC_062569.1_RagTag', 'C2',
  'NC_062570.1_RagTag', 'D1',
  'NC_062571.1_RagTag', 'D2',
  'NC_062572.1_RagTag', 'D3',
  'NC_062573.1_RagTag', 'D4',
  'NC_062574.1_RagTag', 'E1',
  'NC_062575.1_RagTag', 'E2',
  'NC_062576.1_RagTag', 'E3',
  'NC_062577.1_RagTag', 'F1',
  'NC_062578.1_RagTag', 'F2',
  'NC_062579.1_RagTag', 'X',
  'NW_025927612.1_RagTag', 'NW_025927612.1',
  'NW_025927619.1_RagTag', 'NW_025927619.1'
)

chr_df_named <- chr_df %>% 
  left_join(scaff_names, by = c("chr" = "scaffold")) %>% 
  ungroup() %>% 
  dplyr::select(-chr) %>% 
  relocate(chr_name, .before = start)

df_id_chr <- df_id %>% 
  left_join(scaff_names, by = c("chr" = "scaffold")) %>% 
  ungroup() %>% 
  dplyr::select(-chr) %>% 
  relocate(chr_name, .after = ctg)

write_tsv(chr_df_named, "pretty_chrs_chromomap.txt",
          col_names = F)
write_tsv(df_id_chr, "pretty_ctgs_chromomap.txt",
          col_names = F)
```

Graph:

```
library(chromoMap)

chromoMap("pretty_chrs_chromomap.txt", "pretty_ctgs_chromomap.txt", segment_annotation = T, n_win.factor = 2, left_margin = 100, data_based_color_map=T, data_type="categorical", export.options = T, data_colors = list(c('#0173b2', '#de8f05', '#029e73', '#d55e00', '#cc78bc', '#ca9161', '#fbafe4', '#949494', '#ece133', '#56b4e9', '#0173b2', '#de8f05')))
```

#### Software versions

| Software | Version | Application |
| --- | --- | --- |
| AGAT | 0.9.1 | GTF conversion |
| Awk | 20200816 | TXT file manipulation |
| BCFtools | 1.15.1 | Alternate pseudohaplotye |
| BEDtools | 23.3.0 | Nearest features |
| BLAST | 2.13.0 | Contamination |
| Blobtools | 3.1.11 | Contamination |
| BUSCO | 5.3.2 | Assembly quality |
| chromoMap | 4.1.1 | Ideogram visualization |
| Dfam | 3.5 | Called by RepeatMasker |
| DSS | 2.43.2 | Differential DNA methylation |
| Flye | 2.9 | Assembly |
| GeMoMa | 1.8 | Gene annotation |
| Guppy | 6.2.1 | Base calling |
| Hmmsearch | 3.1 | Called by BUSCO |
| IQ-TREE | 1.6.12 | Mitchochondrial phylogeny |
| JBrowse 2 | 1.7.7 | Alignment visualization |
| Kraken2 | 2.1.2 | Contamination |
| Longphase | 1.2 | Haplotagging for Methylartist |
| Medaka | 1.6.0 | Polishing |
| Merqury | 1.3 | Assembly quality |
| Metaeuk | 5.34c21f2 | Called by BUSCO |
| Methylartist | 1.2.3 | DMR visualization |
| Minimap2 | 2.22-r1101 | Alignment |
| Modbam2bed | 0.6.2 | DNA methylation file handling |
| Mosdepth | 0.3.3 | Assembly quality |
| MUMmer4 (nucmer) | 4.0.0cr1 | Pairwise genome alignment |
| MUSCLE | 5.1 | Mitochondrial multi-alignment |
| Nanoq | 0.9.0 | Sequencing statistics |
| NextDenovo | 2.5.0 | Assembly |
| NextPolish | 1.4 | Polishing |
| Pavian | 1.0 | Contamination |
| PEPPER-Margin-DeepVariant | 0.8.0 | Diploid variant calling |
| Purge Haplotigs | 1.1.2 | Assembly cleanup |
| Quickmerge | 0.3 | Assembly merging |
| Racon | 1.4.3 | Polishing |
| Raven | 1.8.1 | Assembly |
| RepeatMasker | 4.1.0 | Repetitive DNA identification |
| SAMtools | 1.15.1 | SAM/BAM handling, coverage data |
| SeqKit | 2.2.0 | Assembly statistics, FASTA handling |
| Seqtk | 1.3 | Read downsampling for mtDNA |
| Shasta | 0.8 | Assembly |
| SMC++ | 1.15.2 | Runs of homozygosity |
| VCFtools | 0.1.17 | VCF file handling |
| Whatshap | 1.4 | Variant statistics |

#### References

Wanner, Nicole, Peter A. Larsen, Adam McLain, and Christopher Faulk.
2021. “The Mitochondrial Genome and Epigenome of the Golden Lion
Tamarin from Fecal DNA Using Nanopore Adaptive Sequencing.”
*BMC Genomics* 22 (1). https://doi.org/10.1186/s12864-021-08046-7.
